# Specific anterior-posterior brain-wide input patterns support specialized visuospatial processing in the mouse retrosplenial cortex

**DOI:** 10.1101/2025.06.24.661247

**Authors:** Yu-Ting Wei, João Couto, Fabian Kloosterman, Vincent Bonin

## Abstract

The retrosplenial cortex (RSC) is a key integrative hub involved in spatial orientation, navigation, and cognitive processes. In rodents, RSC neurons carry rich sensory and navigational signals and are interconnected with sensory, motor, thalamic, and hippocampal circuits—supporting multimodal integration. However, the circuitry that supports this integration remain unclear. Here, we combined 2-photon calcium imaging in navigating mice with brain-wide retrograde tracing to investigate how visual and positional information are represented and distributed across RSC subregions. We found a clear anterior–posterior gradient: anterior RSC neurons exhibited sharper, more reliable position tuning and preferred fast-moving visual stimuli, while posterior RSC neurons showed broader tuning and preferential responses to slower motion. These functional differences were paralleled by distinct patterns of long-range input: anterior RSC received denser projections from motor, parietal, and hippocampal-associated areas—regions implicated in position encoding—whereas posterior RSC was more strongly innervated by visual cortices. Our findings reveal that the RSC contains functionally and anatomically distinct subregions specialized for processing different visuospatial features, suggesting a modular organization that supports integration of contextual and sensory information during navigation.

## INTRODUCTION

The retrosplenial cortex (RSC) plays a central role in diverse cognitive and emotional processes, acting as a hub for integrating sensory, mnemonic, and contextual information to support navigation and learning (Vann et al., 2009; Auger et al., 2012; Epstein et al., 2017; Alexander et al., 2023; Saleem and Busse, 2023). Through its widespread connections with sensory, motor, thalamic, and hippocampal circuits (Yamawaki et al., 2016; Haugland et al., 2019; Brennan et al., 2021; Morimoto et al., 2021), the rodent RSC is well-positioned to transform multimodal information into behaviorally relevant representations that guide navigation and decision-making (Miller et al., 2014; Vedder et al., 2017; Sun et al., 2021; Hattori and Komiyama, 2022). However, the organizational principles that govern multimodal information flows within RSC remain poorly understood.

During spatial navigation, the brain continuously updates the animal’s position and orientation by integrating incoming sensory information to maintain alignment between internal representations and the external environment (Alexander and Nitz, 2015). In rodents, RSC neurons encode a variety of spatially relevant signals, including landmark cues (Vedder et al., 2017), reward locations (Miller et al., 2019), head direction (Jacob et al., 2017; Fallahnezhad et al., 2023; Sit and Goard, 2023), and egocentric boundaries (Alexander et al., 2020; van Wijngaarden et al., 2020; Cheng et al., 2024). These representations are essential for spatial learning and memory, and lesions or inactivation of the RSC result in pronounced navigational impairments (Vann and Aggleton, 2005; Hindley et al., 2014; Fischer et al., 2020).

Vision plays a key role by providing information about environmental landmarks that anchor internal spatial maps. The RSC, through its reciprocal connections with both visual and hippocampal systems, is uniquely positioned to support this alignment. RSC neurons exhibit both visually evoked responses and spatial selective firing (Sugar et al., 2011; Murakami et al., 2015; Mao et al., 2017; Nitzan et al., 2020; Powell et al., 2020; Morimoto et al., 2021). The convergence of visual sensitivity and spatial tuning suggests that the RSC functions as an interface between sensory perception and spatial memory, enabling animals to maintain accurate orientation and effectively coordinate navigation.

Anatomically, the rodent RSC includes ventrally and dorsally located granular and agranular regions, each with distinct lamination and connectivity profiles. Previous studies have reported differences between these subdivisions in memory-related functions, particularly contextual fear (Wang et al., 2019; Yamawaki et al., 2019a; Tsai et al., 2022). Functional differences have also been observed along the anterior-posterior axis, with the posterior RSC more engaged in the encoding and retrieval of fear-related memories (Trask and Helmstetter, 2022).

Despite growing evidence of subregional specialization, less is known about how anterior and posterior RSC contribute to spatial navigation and visuospatial processing. Some evidence suggests hippocampal and visual inputs may be topographically organized along this axis (Niell and Stryker, 2008; Jin and Glickfeld, 2020; Goldbach et al., 2021; Han et al., 2022), raising the possibility that functional gradients exist within RSC.

In this study, we combined in vivo 2-photon calcium imaging and brain-wide retrograde circuit mapping to investigate how cellular activity and long-range input patterns support the encoding of visual and spatial information in the mouse RSC. By targeting excitatory neurons in anterior and posterior RSC, we uncovered subregion-specific specializations in spatial and visual processing. Anterior RSC neurons exhibited sharper and more reliable position tuning, along with stronger responses to high-speed visual motion. In contrast, posterior RSC neurons show broader and weaker position tuning and greater visual responsiveness to slow visual motion. The functional findings are paralleled by distinct anatomical connectivity: anterior RSC receives denser input from motor, parietal, and hippocampal-associated areas, while posterior RSC is more strongly innervated by visual cortices. Together, these findings reveal a topographically organized functional architecture within the RSC, suggesting that anterior and posterior subregions contribute complementary information during visuospatial integration. Future causal studies will be needed to determine the specific contributions of each subregion.

## RESULTS

To investigate how spatial and visual information is functionally organized across the retrosplenial cortex (RSC), we combined large-scale 2-photon calcium imaging with brain-wide retrograde circuit mapping in awake, head-fixed mice. Our goal was to characterize how cellular-level activity and long-range input patterns vary along the caudal surface of RSC. For the functional cellular imaging, we targeted superficial layers of anterior and posterior RSC in mice expressing GCaMP6s (Thy1-GCaMP6s, n= 4 mice, and CaMKII-tTA × TRE-GCaMP6s, n = 4 mice) and recorded neuronal activity during head-fixed locomotion and visual stimulation (Figures 1A–1C). 2-photon imaging was performed in RSPd within the agranular layer of the RSC at depths of 60– 360 µm below the pial surface (Figure 1C; STAR Methods). Mice ran on a linear treadmill with discrete position cues and a fixed reward location to elicit reliable position-related activity (Royer et al., 2012; Mao et al., 2017; Lande et al., 2023). To measure sensory responses, we presented full-field drifting gratings interleaved with gray screen epochs. This experimental design allowed us to assess functional specialization in spatial and visual coding across RSC subregions and to relate these patterns to differences in anatomical connectivity. Recordings were distributed across the anterior-posterior axis: anterior fields of view (FOVs) ranged from –1.04 mm to –2.38 mm relative to Bregma, and posterior FOVs from –2.32 mm to –4.17 mm. Each session began with a 15-minute gray screen period to assess position-related activity during locomotion, followed by a 30-minute block of full-field visual stimulation interleaved with gray screen epochs to measure visually evoked responses (Table S1).

**Figure 1.**
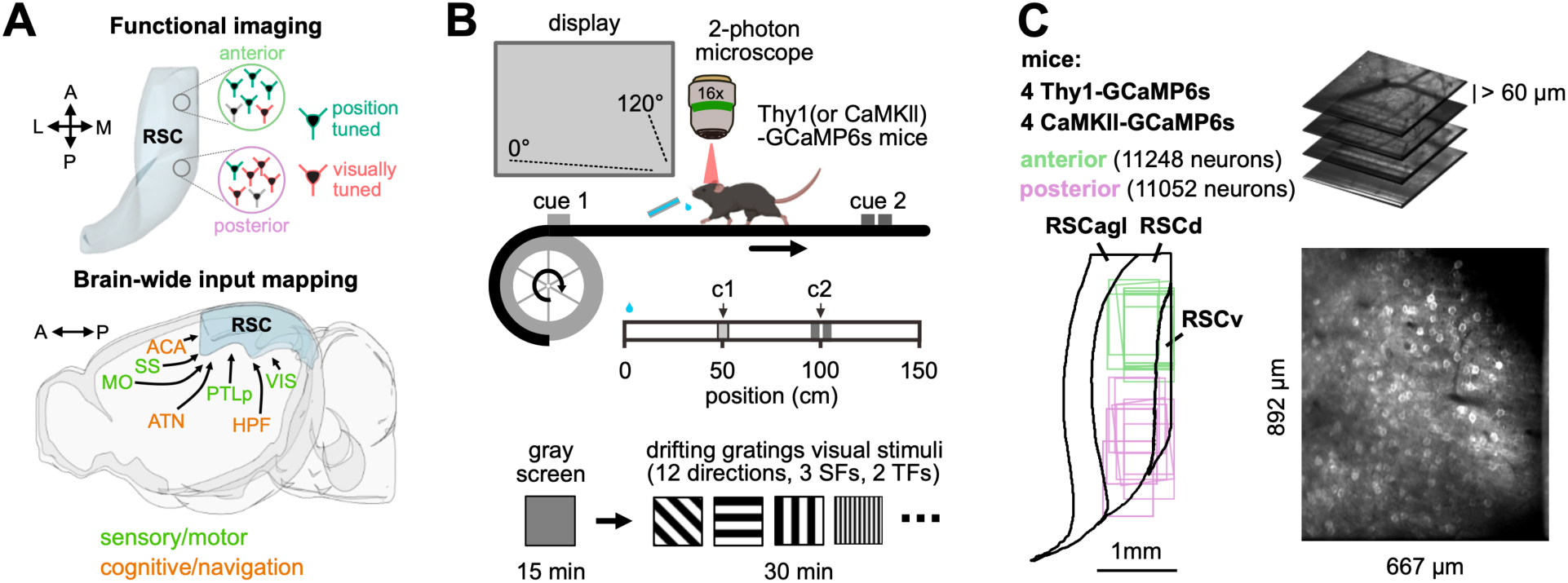
Experimental paradigm for studying functional gradients and long-range connectivity in mouse retrosplenial cortex. (A) Schematic illustration of the experimental framework investigating functional and anatomical specialization along the anterior-posterior axis of the retrosplenial cortex (RSC). Anatomical orientations are labeled: anterior (A), posterior (P), lateral (L), and medial (M). (B) Top: Head-fixed 2-photon imaging setup. Mice ran on a linear treadmill with two position cues (c1 and c2, at 50 cm and 100 cm) and received a water reward at a fixed location (0 cm) on each lap. A monitor was placed 18 cm from the right eye, covering 120 by 80 degrees of the right visual field. Bottom: Each session included a 15-minute gray screen period followed by a 30-minute visual stimulation block with randomized presentations of full-field drifting gratings (12 directions, 3 spatial frequencies, and 2 temporal frequencies). (C) Locations of imaged fields of view (FOVs) in the dorsal cortical surface of the RSC. Each FOV measured 892 × 667 µm. In a subset of experiments, volumetric scanning was performed using an electrically tunable lens (see Table S1).

### Position coding is more robust in anterior than posterior RSC

We first assessed how position-related activity is distributed across the RSC (Mao et al., 2017). Position-selective neurons were identified based on occupancy-normalized, deconvolved calcium activity and a shuffle-based significance test (Figure 2A, left; Figure S1; STAR Methods). A significantly larger fraction of neurons in the anterior RSC exhibited position tuning compared to the posterior RSC (Figure 2A; anterior vs. posterior: 45 ± 5 % vs. 21 ± 3 %; mean ± SEM; p = 0.0017; Mann–Whitney U test; n = 10 sessions from 8 animals). Anterior RSC neurons also showed narrower tuning fields, higher spatial information content, and greater trial-to-trial reliability (Figures 2B–2F; anterior vs. posterior: tuning field width in cm; 45 ± 1.7 vs. 64.3 ± 4.2; p = 0.002; spatial information in bits/event, 0.28 ± 0.02 vs. 0.18 ± 0.01; p < 0.001; median trial-to-trial correlation, 0.17 ± 0.02 vs. 0.11 ± 0.01; p = 0.017; mean ± SEM; Mann–Whitney U test; n = 10 sessions). Spatial information computed from average tuning curves varied systematically along the anterior-posterior axis (Figure 2C). The number of position fields per neuron showed only modest differences between regions (Figure 2G; anterior vs. posterior: 1 field, 56.4 ± 2.4% vs. 68.9 ± 6%; p = 0.1; 2 fields, 37.1 ± 1.7% vs. 27.9 ± 4.8%; p = 0.21; 3 fields, 6.4 ± 0.9% vs. 3.2 ± 1.9%; p = 0.006; mean ± SEM in percentage of position-tuned neurons; Mann–Whitney U test; n = 10 sessions), suggesting that the main distinction lies in tuning precision (narrower tuning field) and reliability. These results indicate that position-related signals are more robustly encoded in anterior RSC compared to posterior RSC.

**Figure 2.**
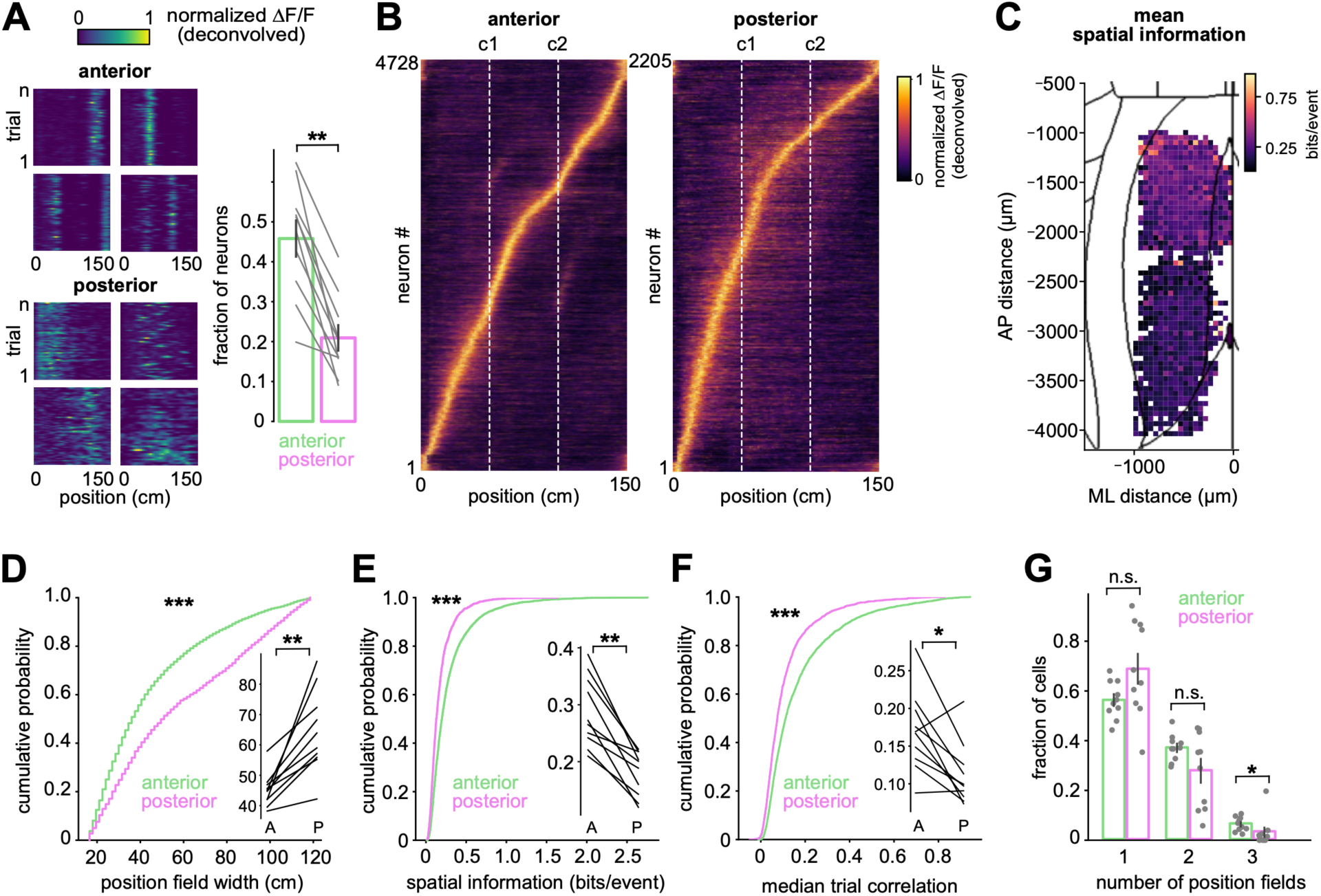
Anterior RSC neurons exhibit enhanced position coding with narrower tuning fields, higher spatial information, and increased reliability. (A) Left: Examples of normalized deconvolved calcium activity (ΔF/F0) from individual position-tuned RSC neurons, plotted as a function of the animal’s position across trials. Right: Proportion of position-tuned neurons in anterior versus posterior RSC. Vertical bars indicate mean ± SEM across sessions; each connected line represents an individual animal-recording session (n = 10 sessions from 8 animals). Mann– Whitney U test; p = 0.0017. (B) Trial-averaged deconvolved ΔF/F0 activity for all position-tuned neurons in anterior and posterior RSC. (C) Spatial distribution of mean spatial information across position-tuned neurons, projected onto the dorsal cortical surface (n = 10 sessions). (D–F) Cumulative distributions comparing (D) tuning field width, (E) spatial information (bits/event), and (F) trial-to-trial correlation coefficients between anterior and posterior RSC neurons. Animal- and recording-wise mean values are shown with connected lines. Statistical comparisons were performed using Kolmogorov–Smirnov (KS) tests on pooled data (4728 vs. 2205 neurons) and Mann–Whitney U tests on per-session averages (n = 10 sessions): KS test vs. Mann–Whitney test; tuning field width, p < 0.001 vs. p = 0.002; spatial information, p < 0.001 vs. p < 0.001; median trial correlation, p < 0.001 vs. p = 0.017. (G) Fraction of neurons with one, two, or three position fields. Vertical bars indicate mean ± SEM across sessions. Mann–Whitney tests: 1 field, p = 0.10; 2 fields, p = 0.21; 3 fields, p = 0.006 (n = 10 sessions).

To determine how these single-neuron differences across RSC translate to population-level coding, we tested how accurately neural ensembles in anterior and posterior RSC could predict the animal’s position. Anterior RSC exhibited more robust activation during locomotion compared to posterior RSC (Figures 3A–3B, top). A Bayesian decoder trained on population ΔF/F_0_ yielded significantly lower decoding errors in anterior RSC (Figure 3A–3B, bottom, 3C and 3E; anterior vs. posterior: 5.2 ± 0.7 cm vs. 10.4 ± 1.2 cm; mean ± SEM; p = 0.0017; Mann–Whitney U test; n = 10 sessions from 8 animals). Additionally, calcium response amplitudes were modestly higher in anterior RSC (Figure 3D; anterior vs. posterior: 110 ± 4.1 vs. 95.9 ± 3.9; mean ± SEM; p = 0.037; Mann–Whitney U test; n = 10 sessions), indicating stronger ensemble engagement in this region. To assess whether the observed regional differences in position coding depend on visual input, we trained the position decoder on neural activity recorded during light conditions and tested it on data recorded in darkness (Figure S2; Table S2; n = 9 sessions from 7 animals). This cross-condition decoding strategy allowed us to evaluate how well position representations in RSC generalize when visual input is removed. Across RSC subregions, decoding error increased significantly in the absence of visual input (Figure S2A–S2B; light vs. dark: 7.8 ± 0.9 cm vs. 15.7 ± 1.6 cm; mean ± SEM; p = 0.0039; Wilcoxon signed-rank test; n = 9 sessions), yet the anterior-posterior gradient in performance remained. Posterior RSC neurons showed a trend toward greater median decoding error (anterior vs. posterior: 13.4 ± 0.7 cm in 6 sessions vs. 20.1 ± 3.8 cm in 3 sessions; mean ± SEM) and reduced position selectivity in the darkness (Figure S2C; anterior vs. posterior: 29.3 ± 5.6% in 6 sessions vs. 9.9 ± 0.5% in 3 sessions; mean ± SEM). While both subregions maintained position-related activity, reliability and response amplitude decreased under dark conditions (Figures S2D–S2F). These findings suggest that although visual input enhances spatial representations, RSC position coding does not rely exclusively on vision. Notably, anterior RSC continued to show stronger and more reliable coding, even in darkness, indicating intrinsic regional differences in contextual processing.

**Figure 3.**
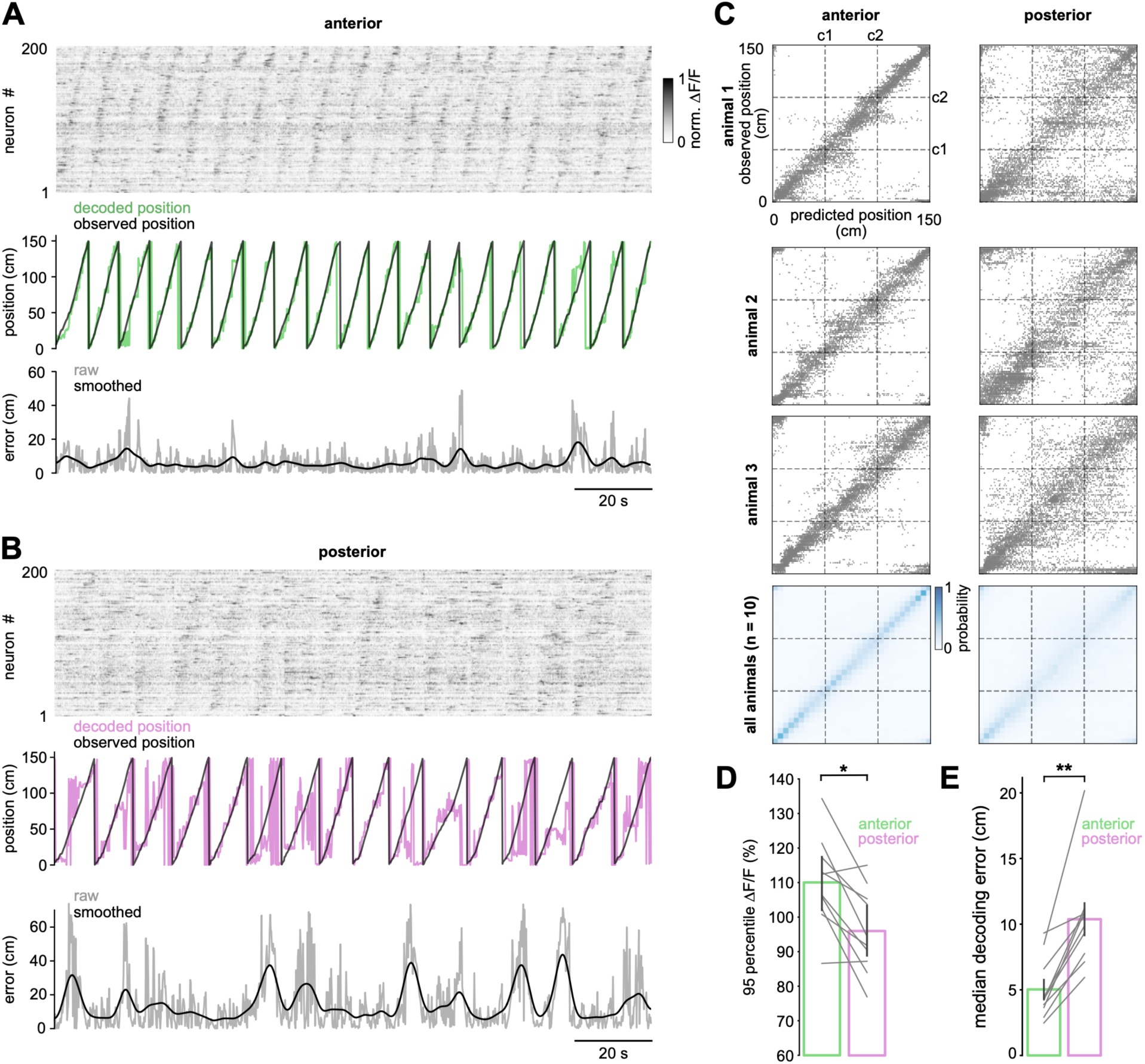
Anterior RSC neurons encode position more accurately than posterior RSC neurons. (A–B) Top: Normalized calcium activity (ΔF/F0) from 200 randomly selected RSC neurons, sorted by position eliciting peak calcium activity. Middle: Observed (black) and decoded (green and magenta) position over time for the same session. Bottom: Decoding error (absolute difference between observed and predicted position), smoothed with a Gaussian kernel (SD = 10 time bins). (C) Top: Confusion matrices showing predicted versus observed position for anterior and posterior RSC neurons from three animals. Bottom: Mean confusion matrix calculated across animals and recordings sessions (n = 10 sessions from 8 animals). Cue locations (c1 and c2) are marked with dotted lines. (D) 95th percentile ΔF/F₀ amplitude in anterior versus posterior RSC. Vertical bars represent mean ± SEM; each connected line indicates an individual animal and recording session. Mann–Whitney U test, p = 0.037. (E) Median decoding error in anterior versus posterior RSC. Vertical bars indicate mean ± SEM across sessions; each connected line indicates an individual animal-recording session. Mann–Whitney U test, p = 0.0017.

### Anterior and posterior RSC differ in visual responsiveness and stimulus preferences

We next examined how visual activity is organized across the RSC. Rodent RSC receives input from both primary and higher visual areas and has been shown to exhibit tuned responses to visual stimuli (Murakami et al., 2015; Zhuang et al., 2017). Widefield calcium imaging revealed a clear retinotopic organization in the posterior RSC, with stronger responses to stimuli presented in the nasal upper visual field. In contrast, anterior RSC showed weaker visual responses and preferentially activation by stimuli in the nasal lower visual field (Figure S3; Supplemental Video S1).

To characterize visual tuning at the cellular level, we recorded responses to repeated presentations of full-field gratings spanning 6 orientations and 12 directions (0°–330°), 3 spatial frequencies (SF: 0.04, 0.08, 0.16 cpd), and 2 temporal frequencies (TF: 1, 4 Hz) (0°–120° azimuth, ± 40° elevation), reflecting the range of visual tuning observed in visual cortex of awake mice (Andermann et al., 2011; Han et al., 2022). Visually responsive neurons were defined as those with a median pairwise trial-to-trial correlation > 0.3 (Figures 4A–4B and Figure S4; STAR Methods). Neurons in both anterior and posterior RSC showed clear visual responses with similar response amplitudes (Figures 4B–4C). However, consistent with the widefield observations, posterior RSC neurons showed greater response reliability and a significantly larger proportion of visually responsive neurons (Figure 4D; anterior vs. posterior: 16.1 ± 2.7% vs. 33.8 ± 3.3%; mean ± SEM; p = 0.0017; Mann–Whitney U test; n = 10 sessions from 8 animals).

**Figure 4.**
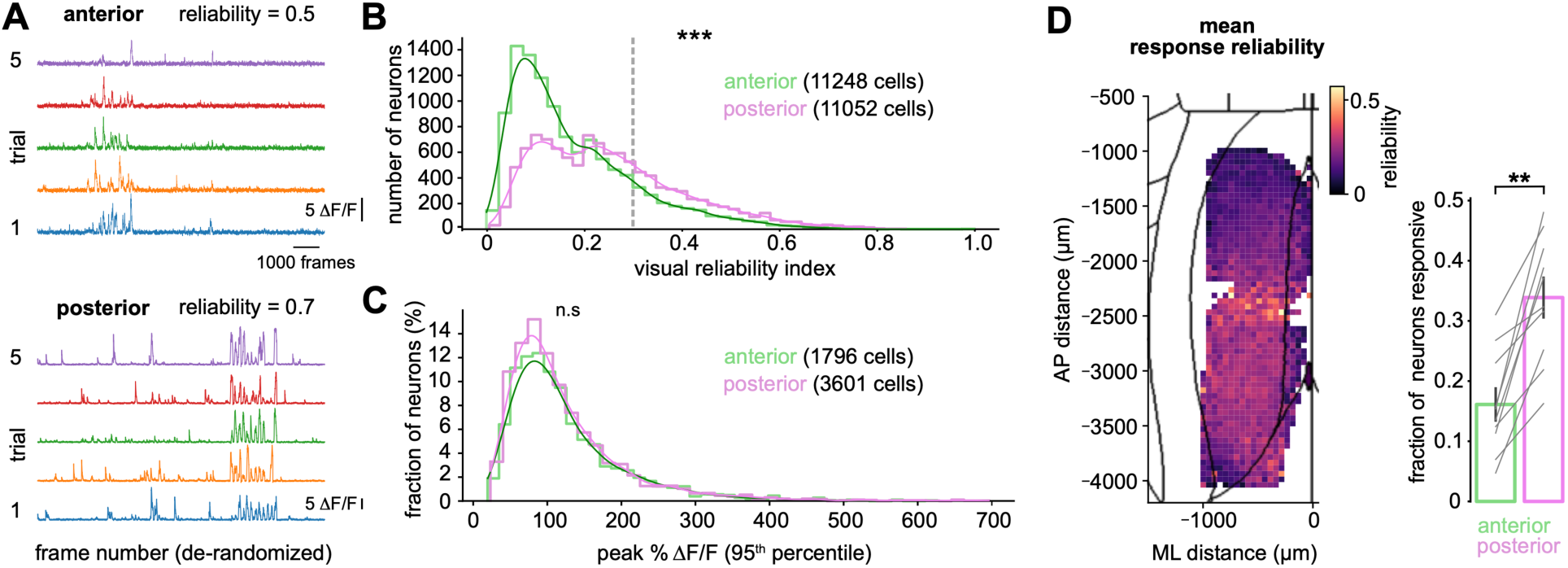
Posterior RSC neurons exhibit greater visual reliability than anterior RSC neurons. (A) Example ΔF/F0 time courses used for visual reliability assessment. Responses to the stimuli were de-randomized and sorted by trial (rows). Reliability index (") was computed as the 75th percentile of trial-to-trial Pearson correlation coefficients. (B) Distribution of visual reliability index for all recorded neurons across anterior and posterior RSC. The dotted line indicates the threshold (0.3) used to classify a neuron as visually responsive. Kolmogorov-Smirnov test; p < 0.001; 11248 (anterior) vs. 11052 (posterior) neurons (n = 10 sessions from 8 animals). (C) Distribution of maximal ΔF/F0 amplitudes (95% percentile across individual trials) for visually responsive neurons. Kolmogorov-Smirnov test; p = 0.08; n = 1796 (anterior) vs. 3601 (posterior) neurons (n = 10 sessions). (D) Left: Spatial map of visual reliability across the dorsal cortical surface. Right: Proportion of visually-responsive neurons in anterior versus posterior RSC. Vertical bars indicate mean ± SEM across sessions. Each connected line represents an individual animal. Mann–Whitney U test; p = 0.0017 (n = 10 sessions).

Further analysis revealed distinct spatial and temporal frequency tuning between RSC subregions. Anterior RSC neurons preferentially responded to low spatial and high temporal frequencies, such as 0.04 cycles per degree (cpd) at 4 Hz (Figure 5A–5B, top), whereas posterior RSC neurons were more responsive to high spatial and low temporal frequencies, such as 0.16 cpd at 1 Hz (Figure 5A–5B, bottom). These preferences were reflected in the proportions of neurons responsive to each SF/TF combination (Figure 5C; anterior vs. posterior: 0.04 cpd at 4 Hz: 28.3 ± 3.5% vs. 5.5 ± 1.1%, p < 0.001; 0.08 cpd at 4 Hz: 32 ± 1.7% vs. 10.4 ± 2.5%, p < 0.001; 0.16 cpd at 4 Hz: 15.4 ± 2.3% vs. 21.1 ± 2.0%, p = 0.18; 0.04 cpd at 1 Hz: 11.6 ± 2.1% vs. 22.8 ± 1.5%, p = 0.002; 0.08 cpd at 1 Hz: 5.9 ± 1.5% vs. 15.2 ± 1.5%, p = 0.0017; 0.16 cpd at 1 Hz: 6.5 ± 1.5% vs. 25.0 ± 2.7%, p < 0.001; mean ± SEM; Mann–Whitney U test; n = 10 sessions from 8 animals). The tuning bias in anterior RSC favors fast, low-resolution stimuli, whereas posterior RSC is tuned for slower, high-resolution visual input. Such specialization parallels the gradient of tuning properties observed along the anterior-posterior dimension of visual cortical areas (Andermann et al., 2011; Han et al., 2022).

**Figure 5.**
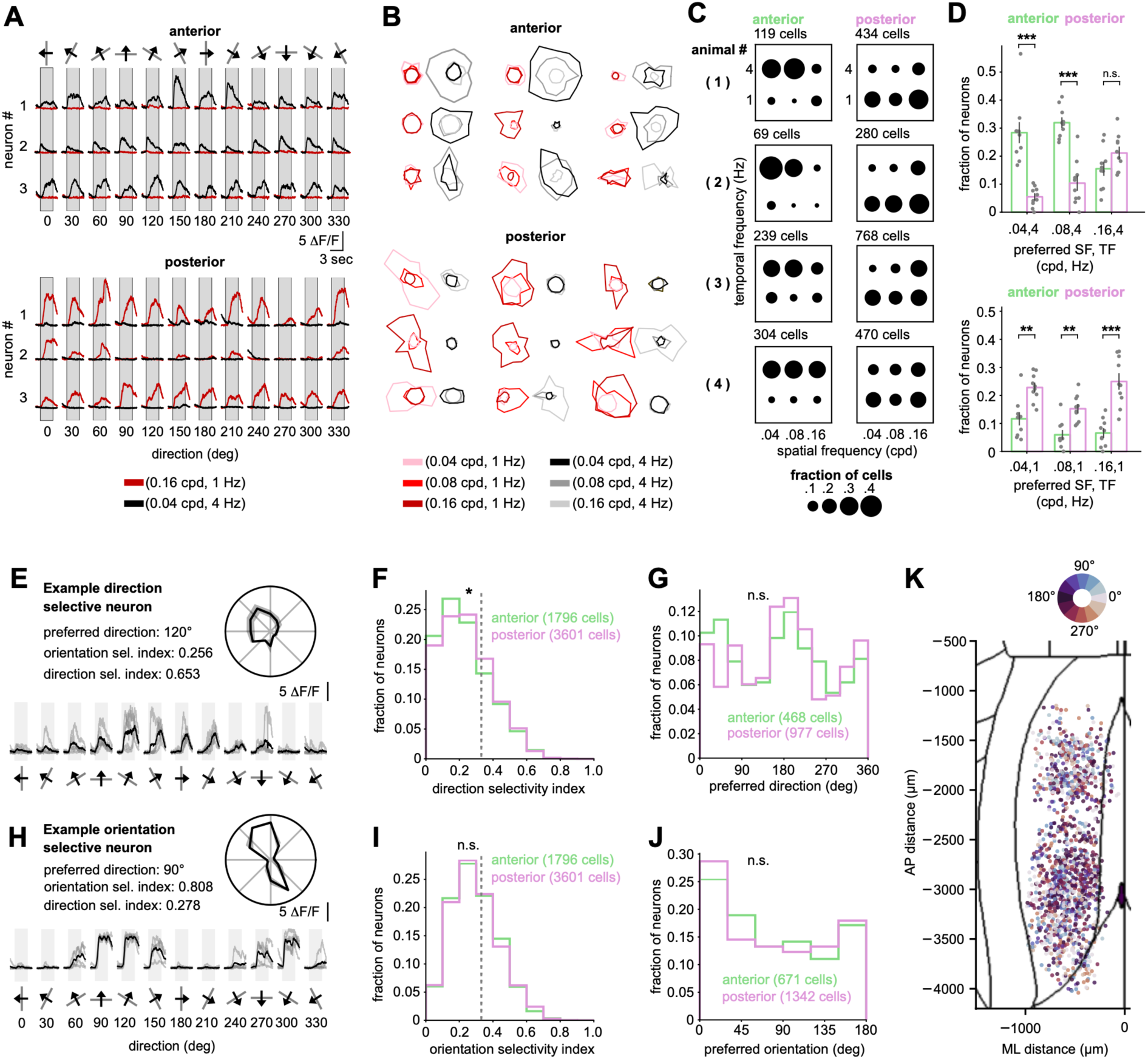
Anterior and posterior RSC visual neurons encode distinct spatial and temporal frequencies. (A) Example calcium traces from visually responsive neurons in anterior and posterior RSC. Each column corresponds to a different stimulus direction (30° increments), and each color corresponds to a distinct spatial and temporal frequency (SF/TF) combination. (B) Examples of visual tuning shown in polar plots. The angle represents stimulus direction; color encodes the spatial and temporal frequency combination. (C) Dot plots from four representative animals showing the proportion of visually responsive neurons tuned to each SF/TF combination. The proportion of cells is shown as surface area. (D) Group-level analysis of SF/TF tuning preferences in anterior and posterior RSC. Vertical bars indicate mean ± SEM across recording sessions. Mann–Whitney U test; p < 0.001 (0.04 cpd, 4 Hz); p < 0.001 (0.08 cpd, 4 Hz); p = 0.18 (0.16 cpd, 4 Hz); p = 0.002 (0.04 cpd, 1 Hz); p = 0.0017 (0.08 cpd, 1 Hz); p < 0.001 (0.16 cpd, 1 Hz); n = 10 sessions from 8 animals. (E) Example direction tuning curve and corresponding polar plot for a direction-selective neuron. Direction selectivity index and orientation selectivity index are shown. (F) Histogram of direction selectivity index values for visually responsive neurons in anterior and posterior RSC. The dotted line (0.33) indicates the classification threshold for direction selectivity. Kolmogorov-Smirnov test; p = 0.012; 1796 (anterior) vs. 3601 (posterior) neurons; n = 10 sessions. (G) Histogram of preferred directions among direction-selective neurons in anterior and posterior RSC. Kolmogorov-Smirnov test; p = 0.064; 468 vs. 977 neurons; n = 10 sessions. (H) Example orientation tuning curve and corresponding polar plot for an orientation-selective neuron. (I) Histogram of orientation selectivity index values for visually responsive neurons. The dotted line (0.33) indicates the classification threshold. Kolmogorov-Smirnov test; p = 0.88; 1796 vs. 3601 neurons; n = 10 sessions. (J) Histogram of preferred orientations for orientation-selective neurons in anterior and posterior RSC. Kolmogorov-Smirnov test; p = 0.74; 671 vs. 1342 neurons; n = 10 sessions. (K) Dorsal cortical map showing the location of individual visually responsive neurons color-coded by preferred direction.

In contrast to spatial and temporal tuning, orientation and direction selectivity showed less pronounced differences along the anterior-posterior RSC axis (Figures 5E–5K). Neurons in both anterior and posterior RSC exhibited orientation- and direction-selective responses. No significant differences were observed in orientation selectivity index (p = 0.88), preferred orientation (p = 0.74), or preferred direction (p = 0.064; Kolmogorov–Smirnov test; n = 10 sessions). The proportions of direction-selective (anterior vs. posterior: 26.1% vs. 27.1%) and orientation-selective neurons (anterior vs. posterior: 37.4% vs. 37.3%) were also comparable between subregions. However, posterior RSC showed a modest but significant increase in direction selectivity index (Figure 5F, p = 0.012, Kolmogorov-Smirnov test; n = 10 sessions), possibly reflecting differences in spatial and temporal preferences (Figure 5C–D). Notably, we observed no evidence of topographic organization in the preferred orientation or direction within either region (Figure 5K).

In summary, visual signals are unevenly distributed across the RSC. Posterior RSC showed more robust and reliable activity, particularly those requiring high spatial resolution, while anterior RSC displayed weaker visual responsiveness but is more strongly modulated by task context. These findings highlight a functional gradient along the anterior-posterior axis of the RSC, pointing to distinct integrative roles.

### Afferent connectivity correlates with RSC representations

The retrosplenial cortex (RSC) integrates input from a wide range of cortical and subcortical areas, including visual cortices, the hippocampal formation, anterior thalamic nuclei, posterior parietal cortex, and sensorimotor areas (Van Groen and Wyss, 2003; Wang and Burkhalter, 2007; Yamawaki et al., 2016, 2019b; Haugland et al., 2019; van Wijngaarden et al., 2020; Brennan et al., 2021; Sit and Goard, 2023; Li et al., 2025). We hypothesized that functional differences along the anterior-posterior RSC axis reflect distinct patterns of long-range input. To test this, we performed brain-wide, cellular-resolution mapping of afferent projections to anterior and posterior RSC using retrograde AAV tracers (Figures 6B–6C; Table S3). Injections were targeted to anterior (–1.5 mm AP) and posterior (–3.2 mm AP) RSC, and labeled cell bodies were registered to the Allen Mouse Brain Atlas (Figures 6D–6E). We quantified both volume-normalized cell density and fraction of inputs across brain regions based on the hierarchical tree structure map (Wang et al., 2020).

**Figure 6.**
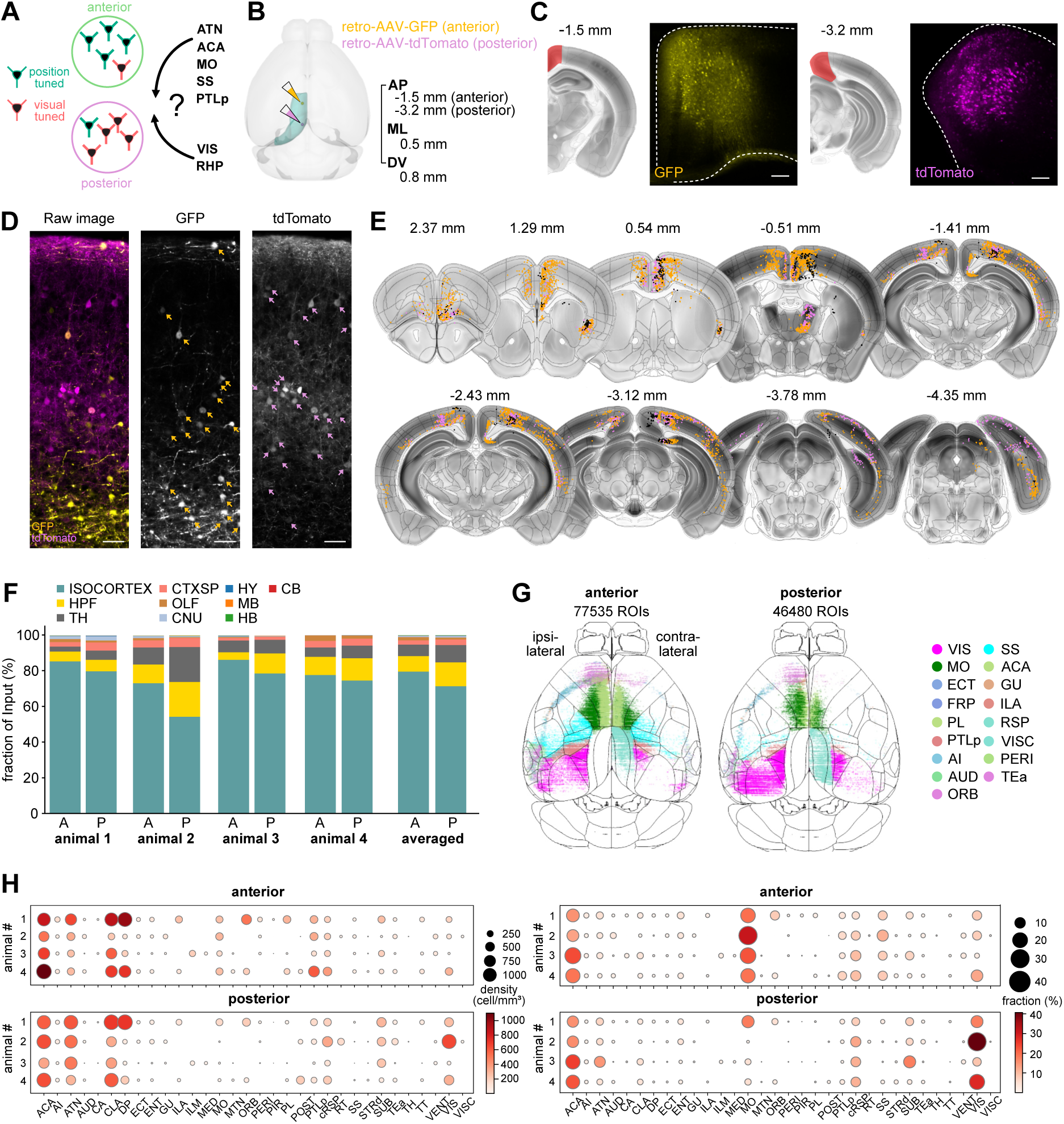
Brain-wide retrograde mapping reveals distinct long-range input sources to anterior and posterior RSC. (A) Schematic of the experimental design for mapping differential afferent connectivity to anterior and posterior RSC using retrograde viral tracing. (B) Retrograde AAV constructs encoding fluorescent proteins were injected into anterior and posterior RSC in wild-type mice to label upstream input neurons. (C) Fluorescent expression at the injection site three weeks post-injection, shown in coronal brain sections. Scale bar, 100 µm. (D) Left: Example stack image showing retrogradely labeled neurons. Middle and Right: Identification of the ROIs expressing GFP and tdTomato from individual fluorescence channels. Scale bar, 50 µm. (E) Example of serial coronal sections showing identified retrogradely labeled neurons. GFP-expressing (gold), tdTomato-expressing (magenta) neurons, and overlapping (gray) neurons are displayed. Coordinates relative to Bregma are indicated above each section. (F) Distribution of labeled neurons across major brain regions for four animals. Abbreviations: HPF, hippocampal formation; TH, thalamus; CTXsp, cortical subplate; OLF, olfactory areas; CNU, cerebral nuclei; HY, hypothalamus; MB, midbrain; HB, hindbrain; CB, cerebellum. (G) Dorsal cortical projection view showing the spatial distribution of labeled input neurons across the isocortex. (H) Brain-wide input distribution by region. Left: volume-normalized cell density. Right: proportion of total labeled input neurons. Dot size and color indicate relative input strength. Abbreviations: ACA, anterior cingulate area; AI, agranular insular area; ATN, anterior group of the dorsal thalamus; AUD, auditory areas; VIS, visual areas; CA, ammon’s horn; CLA, claustrum; DP, dorsal peduncular area; ECT, ectorhinal area; ENT, entorhinal area; GU, gustatory areas; ILA, infralimbic area; ILM, intralaminar nuclei of the dorsal thalamus; MED, medial group of the dorsal thalamus; MO, somatomotor areas; MTN, midline group of the dorsal thalamus; ORB, orbital area; PERI, perirhinal area; PIR, piriform area; PL, prelimbic area; POST, postsubiculum; PTLp, posterior parietal association areas; RSP, retrosplenial area; RT, reticular nucleus of the thalamus; SS, somatosensory areas; STRd, striatum dorsal region; SUB, subiculum; TEa, temporal association areas; VENT, ventral group of the dorsal thalamus; VISC, visceral area.

Most RSC-projecting neurons originated from the cortex, comprising 80.7% of inputs to anterior RSC and 71.9% to posterior RSC (Figure 6F–6G). The hippocampal formation and thalamus also contributed substantially (HPF: 7.6% and 12.4%, TH: 6.1% and 9.9%, anterior vs. posterior. Figure 6F–6H; mean across n = 4 animals).

Major cortical sources of input included the anterior cingulate area (ACA), somatomotor cortex (MO), somatosensory cortex (SS), posterior parietal cortex (PTLp), and visual areas (VIS) (Figure 6H). Consistent with prior work showing position-related input from intracortical pathways to RSC (Gianatti et al., 2023), ACA projected similarly to both subregions (Figure 7A, anterior vs. posterior; 17.7 ± 2.3% vs. 19.8 ± 2.4%; mean ± SEM in fraction of inputs; n = 4 animals), while MO, SS, and PTLp preferentially targeted anterior RSC (Figures 7B–7D, anterior vs. posterior; MO: 23.1 ± 2.7% vs. 7.1 ± 3.8%, SS: 8.7 ± 1% vs. 1.2 ± 0.5%, PTLp: 4.9 ± 0.9% vs. 2.1 ± 0.5%). In contrast, posterior RSC received stronger input from visual areas overall (Figure 7E; anterior vs. posterior: 7.6 ± 2.1% vs. 23.7 ± 6.9%; mean ± SEM in fraction of total; n = 4 animals). Within visual cortical areas (VIS), anterior RSC was more heavily innervated by VISam and VISpor (VISam: 33.6 ± 2.6% vs. 28.4 ± 6.6%; VISpor: 16.2 ± 7.2% vs. 3.8 ± 1.2%; mean ± SEM in fraction of VIS; n = 4 animals), whereas posterior RSC receive more input from VISp and VISpm (anterior vs. posterior: 17.2 ± 3.8% vs. 26 ± 0.6%; mean ± SEM in fraction of VIS inputs; n = 4 animals). This gradient in visual inputs mirrors previously described specialization within visual cortical projections to associative areas and closely matches the tuning properties observed in anterior and posterior RSC neurons—where anterior RSC preferentially receives input from areas tuned to fast, low-resolution stimuli, and posterior RSC is targeted by regions encoding slower, high-resolution features (Marshel et al., 2011; de Vries et al., 2020; Han et al., 2022).

**Figure 7.**
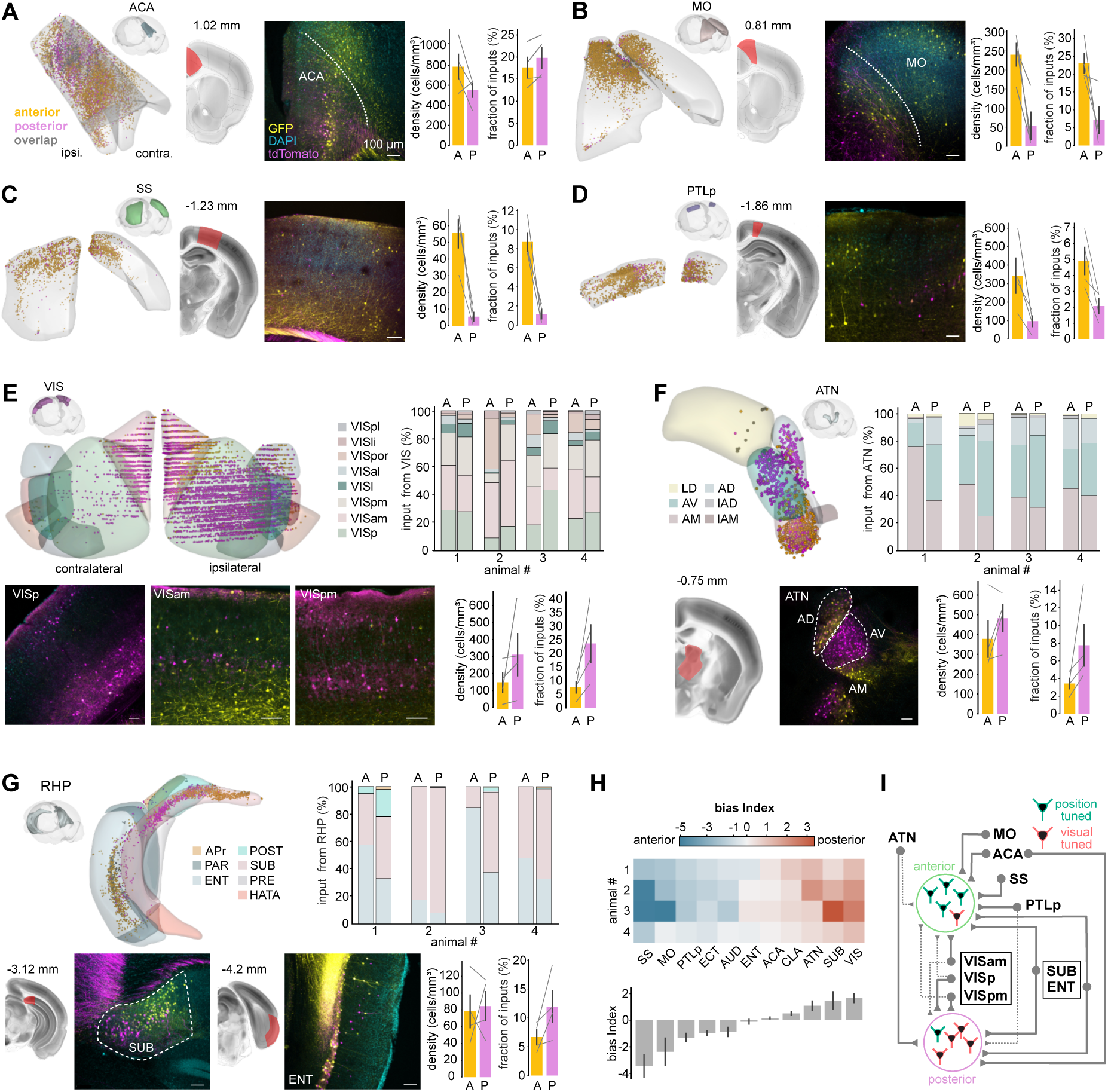
Anterior and posterior RSC neurons receive distinct afferent inputs from sensorimotor, visual, thalamic, and hippocampal regions. (A–D) Retrogradely labeled neurons in ACA, MO, SS and PTLp, respectively. First column: 3D reconstruction of the brain area with labeled neurons in one animal. Second and third columns: confocal image and corresponding coronal slice. Fourth and fifth columns: quantification of volume-normalized density and proportion of labeled neurons across animals. Vertical bars indicate mean ± SEM (n = 4 animals). (E–G) Same as (A–D) but showing the labeled neurons and reconstruction in visual areas (VIS), anterior dorsal thalamus (ATN) and retrohippocampal area (RHP). Dot plots indicate the proportion of each subregion within the brain area across animals. Abbreviations: VISal, anterolateral visual area; VISam, anteromedial visual area; VISl, anterolateral visual area; VISpm, posteromedial visual area; VISli, laterointermediate area; VISp, primary visual area; VISpor, postrhinal area; VISpl, posterolateral visual area. AM, anteromedial nucleus; AV, anteroventral nucleus; AD, anterodorsal nucleus; LD, lateral dorsal nucleus; IAD, interanterodorsal nucleus; IAM, interanteromedial nucleus. ENT, entorhinal area; SUB, subiculum; POST, postsubiculum; APr, area prostriata, PRE, presubiculum; HATA, hippocampo-amygdalar transition area, PAR, Parasubiculum. (H) Top: Bias index for the selected brain areas for each animal. Bottom: Value were shown in mean ± SEM. (I) Summary diagram integrating physiological and anatomical findings.

The thalamus contributes substantial input, primarily from the anterior dorsal thalamic nuclei (ATN), which make up over 70% of thalamic projections. As a key node in the head-direction system (Jankowski et al., 2015), the ATN differentially targeted RSC subregions: posterior RSC received more input from the anteroventral (AV) nucleus (Figure 7F; 30 ± 4.6% vs. 46.7 ± 4.1%), while anterior RSC was more strongly innervated by the anteromedial (AM) nucleus (52 ± 8.2% vs. 33.2 ± 3.3%; mean ± SEM in fraction of ATN; n = 4 animals). Overall, posterior RSC received a greater share of total ATN input (anterior vs. posterior: 3.4 ± 0.6% vs. 7.8 ± 2.3%; mean ± SEM in fraction of total; n = 4 animals).

We further investigated hippocampal inputs, focusing on the retrohippocampal region (RHP), which comprised over 90% of hippocampal afferents to the RSC (Figure 7G). Within the RHP, more than 95% of labeled neurons were located in the subiculum (SUB) and entorhinal cortex (ENT). While posterior RSC received more input from the RHP overall, anterior RSC was more strongly innervated by ENT (anterior vs. posterior: 51.7 ± 13.8% vs. 27.7 ± 6.7%; n = 4 animals). SUB inputs showed a dorsal–ventral gradient: dorsal SUB targeted anterior RSC, whereas ventral SUB projected to posterior RSC. This organization aligns with known functional specialization in the hippocampus, where dorsal SUB is linked to spatial processing—consistent with our findings that anterior RSC more reliably encodes position.

In summary, these results reveal distinct patterns of afferent inputs along the anterior-posterior axis of the RSC (Figure 7H), aligning with functional specialization in spatial and visual processing (Figure 7I).

## DISCUSSION

Using *in vivo* 2-photon calcium imaging and brain-wide circuit mapping, we investigated how neuronal activity and long-range inputs support the encoding of multimodal information in the mouse retrosplenial cortex (RSC). By targeting excitatory neurons in anterior and posterior RSC, we identified distinct subregional specializations in spatial and visual processing. During head-fixed locomotion, anterior RSC neurons exhibited sharper and more reliable position tuning, while posterior RSC neurons showed broader and weaker responses. In response to visual stimulation, anterior RSC neurons preferred fast-moving, low spatial frequency stimuli, whereas posterior RSC neurons were more responsive to slowly-varying, high-resolution visual motion. These functional differences were paralleled by differential input patterns: anterior RSC received denser projections from motor, parietal, and hippocampal-associated areas, while posterior RSC was more strongly innervated by visual cortices. Together, these findings reveal a functional and anatomical gradient along the RSC’s anterior-posterior axis, suggesting that its subregions make complementary contributions to visuospatial integration and navigation.

### Visuospatial coding emerges along an anterior-posterior gradient in RSC

Studies have primarily classified the RSC based on cytoarchitecture–dividing it into granular and agranular subregions, including dorsal/ventral agranular/granular, and lateral agranular zones (RSPd, RSPv, and RSPagl). Lesion and genetic studies have implicated these subdivisions in contextual fear and spatial memory (Lukoyanov and Lukoyanova, 2006; Tsai et al., 2022) and spatial memory (Vann and Aggleton, 2005). However, most physiological studies have focused on limited subregions, and a full functional comparison across the RSC has been lacking. While recent reports highlight differential encoding between anterior and posterior RSC, they have largely focused on object recognition or contextual fear (de Landeta et al., 2020; Trask et al., 2021; Trask and Helmstetter, 2022), leaving open questions about their respective roles in spatial navigation. Our findings fill this gap by demonstrating region-specific differences in spatial and visual coding, suggesting that anterior and posterior RSC support distinct but complementary components of navigational computation. This functional dissociation provides a framework for future studies to investigate how these subregions contribute to visuospatial integration and memory formation.

### The source of spatial information in the RSC

Recent work has highlighted the diverse cortical and subcortical pathways that convey spatial information to the RSC (Gianatti et al., 2023). Notably, projections from the secondary motor and posterior parietal cortices carry strong positional signals, aligning with our anatomical tracing, which revealed dense inputs to anterior RSC—where position-related activity is most robust. Lesions to the hippocampus or disruption of parahippocampal circuits impair spatial tuning in RSC neurons (Mao et al., 2018; van Wijngaarden et al., 2020), supporting a critical role for hippocampal-derived input in spatial coding in RSC.

Consistent with previous findings (Yamawaki et al., 2019a), we observed sparse direct input from hippocampal CA fields to the RSC. This may partly reflect the injection site locations, which likely missed the ventral portion of granular RSC—the primary recipient of direct CA1 input (Li et al., 2025). Within retrohippocampal regions, the dorsal subiculum stood out as a key hub, projecting preferentially to the anterior RSC (Figure 7G). This structure encodes geometric features such as corners and curvatures (Sun et al., 2024) and integrates CA1 inputs to reconstruct environmental layouts. Moreover, gene expression and connectivity profiles vary along the subiculum’s dorsal– ventral axis (Cembrowski et al., 2018; Ding et al., 2020), suggesting functional specialization across its subdivisions. Together, these findings support a model in which the subiculum serves as a major relay for hippocampal spatial signals to the RSC, with its topographic outputs potentially contributing to functional differentiation along the anterior-posterior axis.

### Implication of visual specializations in the RSC

Our widefield retinotopic mapping revealed a gradient of visual responsiveness across the RSC, extending from the posterolateral to the anteromedial axis during stimulation of the upper to lower nasal visual fields. This gradient suggests that enhanced position-related activity in anterior RSC may be influenced by visual perception of tactile cues located in the lower nasal field. To test the role of vision in driving spatial signals, we recorded neural activity in complete darkness (Figure S2). While the absence of visual input reduced the reliability of position tuning in both anterior and posterior RSC, sequential activity patterns were preserved. This indicates that position coding in anterior RSC is not solely dependent on visual cues but instead reflects the integration of multisensory and motor-related inputs.

We also observed distinct spatiotemporal tuning preferences between RSC subregions: anterior RSC responded more to high temporal, low spatial frequency stimuli, whereas posterior RSC was tuned to low temporal, high spatial frequencies. Similar tuning gradients have been reported in medial higher visual areas such as VISam and VISpm (Han et al., 2022), suggesting these areas differentially target anterior and posterior RSC, respectively, providing a likely anatomical substrate for the observed specialization. These findings support the idea that the RSC is not a homogeneous visual processing area but instead exhibits region-specific visual integration, potentially enabling distinct visuospatial computations across its subregions.

### Thalamic contributions to heading and visual Integration in the RSC

Our anatomical data revealed strong projections from the anterior dorsal thalamic nuclei (ATN) to the RSC, particularly from the anteromedial (AM), anteroventral (AV), and anterodorsal (AD) subregions. These thalamic areas are implicated in head-direction coding, spatial working memory, and hippocampal–cortical communication (Jankowski et al., 2013; Roy et al., 2022; LaChance and Taube, 2024). We observed differential targeting of ATN subregions along the RSC’s anteroposterior axis, with AM and AV projecting preferentially to distinct RSC regions— suggesting functional segregation within the thalamocortical pathway. Although the specific roles of each ATN subdivision remain unclear, this topography may reflect parallel processing streams within RSC circuits. The convergence of thalamic and visual inputs likely contributes to the coregistration of head direction and visual landmarks (Alexander et al., 2020; Sit and Goard, 2023), anchoring allocentric representations and supporting stable navigation in changing environments.

### The role of RSC in visuospatial integration and navigation

An expanding body of research highlights the close interaction between the spatial and visual systems in the brain. Visual landmark plays a crucial role in stabilizing spatial representations within the hippocampal formation, influencing the firing properties and stability of place cells and grid cells (Scaplen et al., 2014; Campbell et al., 2018; Bourboulou et al., 2019). Conversely, spatial context and locomotion modulate visual responses in the primary visual cortex and higher visual areas (Saleem et al., 2018; Mika Diamanti et al., 2021), suggesting dynamic interactions between visual and navigational systems. Within this broader network, the RSC emerged as a central hub that integrates visual cues with spatial and self-motion information and provides feedback projection to visual areas such as V1, potentially modulating their spatial representations.

In this study, we focused on mapping upstream inputs to the RSC and identified dense projections from visual, motor, thalamic, and parahippocampal regions. While we did not trace reciprocal outputs, prior studies have shown that the RSC is embedded in bidirectional circuits with both visual and hippocampal systems (Sugar et al., 2011; Makino and Komiyama, 2015; Haugland et al., 2019; Morimoto et al., 2021; Li et al., 2025). These recurrent loops suggest that the RSC is well-positioned to support both bottom-up sensory integration and top-down modulation during spatial behavior. In addition, intrinsic RSC connectivity may facilitate internal coordination between landmark and position signals (Shibata et al., 2009). While previous works have emphasized the RSC’s role in landmark encoding (Auger et al., 2012; Vedder et al., 2017; Fischer et al., 2020), the specific circuits underlying landmark-based memory consolidation remain poorly understood. Future work combining visually guided navigation tasks with pathway-specific perturbations will be essential to dissect the distinct contributions of RSC inputs and outputs in multisensory spatial cognition.

## METHODS

## RESOURCES TABLE

| RESOURCE | SOURCE | IDENTIFIER |
| --- | --- | --- |
| **Mouse line** |  |  |
| Wild type | KU Leuven | RRID:IMSR_JAX:000664 |
| Thy-1 GCaMP6 line | Jackson Laboratories | RRID:IMSR_JAX:024275 |
| CaMKII GCaMP6 line | Jackson Laboratories | RRID:IMSR_JAX:024742;<br>RRID:IMSR_JAX:007004 |
| **Chemical and virus** |  |  |
| pAAV-CAG-GFP | addgene | # 37825-AAVrg |
| pAAV-CAG-tdTomato | addgene | # 59462-AAVrg |
| pAAV-hSyn-EGFP | addgene | # 50465-AAVrg |
| pAAV-hSyn-mCherry | addgene | # 114472-AAVrg |
| Vetergesic | Ceva Santé Animale | CNK:2623-627 |
| Cefazolin | Sandoz | CNK:2218-204 |
| Ketamine | Dechra | CNK:3120-060 |
| Xylazine | VMD | CNK:1339001 |
| Phosphate-buffered saline | Invitrogen | # AM9625 |
| Histofix | Carl Roth | # P087.5 |
| Agarose | Sigma | # A9793-100G |
| Mounting medium with DAPI | VECTASHIELD | # H-2000 |
| UV Curing Optical Adhesives | Thorlabs | # NOA61 |
| **Software and algorithms** |  |  |
| Fiji | NIH | RRID:SCR_002285 |
| MATLAB R2017-R2020 | Mathworks | https://www.mathworks.com/ |
| SHARP-Track | Shamash et al., 2018 | https://github.com/cortex-lab/allenCCF |
| neuralib | Open Source | https://github.com/ytsimon2004/neuralib |
| rscvp | Open Source | https://github.com/ytsimon2004/rscvp |
| posdc | Open Source | https://github.com/ytsimon2004/posdc |
| brainrender | Claudi et al., 2021 | https://github.com/brainglobe/brainrender |
| Suite2p | https://doi.org/10.1101/061507 | https://github.com/mouseland/suite2p |

## RESOURCE AVAILABILITY

### Lead contact

Further information and requests for resources and reagents should be directed to and will be fulfilled by the lead contact, Vincent Bonin.

### Materials availability

This study did not generate new unique reagents.

### Data and code availability

- 2-photon imaging data and retrogradely labelled data have been deposited at zenodo and are publicly available as of the date of publication. DOIs are listed in the key resources.
- All original code has been deposited on Github and is publicly available as of the date of publication. DOIs are listed in the key resources table.
- Any additional information required to reanalyze the data reported in this paper is available from the lead contact upon request.

## EXPERIMENTAL MODEL AND SUBJECT DETAILS

### Animals

All experimental procedures were approved by the Animal Ethics Committee of KU Leuven. For chronic cellular calcium imaging (see Tables S1 and S2), we used either C57BL/6J-Tg(Thy1-GCaMP6s)GP4.12Dkim/J mice or CaMKII-tTA x TRE-GCaMP6s (line G6s2) mice. Wild-type C57BL/6J mice were used for retrograde tracing experiments (Table S3). All animals were between 3 and 8 months old at the time of the experiments. Mice were single-housed in an enriched environment consisting of cotton bedding, wooden blocks, and a running wheel, under a 12 h light/dark cycle, with room temperature maintained at 19–21°C and relative humidity ranging from 30% to 70%.

## METHOD DETAILS

### Cranial window surgery

To enable chronic cellular calcium imaging, mice were anesthetized with isoflurane (2.5–3% for induction, 1–1.25% for surgery). A custom-made titanium frame was attached to the skull, and a cranial glass window (4 or 5 mm, 2–3 layers) was implanted over the left hemisphere over theRSC (Goldey et al., 2014). Postoperative care included administration of Buprenex (2 mg/kg) and Cefazolin (5 mg/kg) every 12 h for two days to manage pain and prevent infection.

### Habituation and behavioral training

Following recovery, mice were gradually habituated to head restraint while positioned on a linear treadmill. To motivate locomotion during the behavioral task, animals were placed on a water restriction schedule, receiving no less than 1 mL of water per day. Body weight was monitored daily to ensure it remained above 80% of their free-feeding weight. Habituation progressed from short initial sessions to durations of up to 1 hour per day. During this period, animals were introduced to the experimental setup without tactile cues. A small water reward (∼2–3 µL) was delivered at a fixed location following each completed lap. Additional rewards were occasionally provided to encourage task-related behaviors, such as licking and running.

### 2-photon calcium imaging

During the imaging experiments, two tactile cues were placed at 50 cm and 100 cm along the treadmill belt to enhance position coding stability in the RSC. Eye movements and facial expressions were simultaneously recorded using an integrated tracking system with AVT Prosilica GC660 and Mako G-030 cameras, respectively.

Cellular calcium imaging was conducted using a resonant scanner -based 2-photon microscope (Neurolabware). In a subset of the experiments, an electrically tunable lens (ETL) was employed to sequentially image across four optical planes, covering a depth of approximately 300 µm from the cortical surface, with an inter-plane spacing of 60–100 µm (see Table S1 and Table S2). For single-plane imaging, data were acquired at ∼30 frames per second, while ETL-based volumetric imaging achieved an effective frame rate of ∼7.5 Hz per plane. The genetically encoded calcium indicator GCaMP6 was excited at 920 nm using a MaiTai DeepSee laser (Spectra Physics / Newport) through a 16x water-immersion objective (NA = 0.8, Nikon). Emitted green light was filtered (510/84 nm, Semrock) and detected using a GaAsP photomultiplier tube (PMT, Hamamatsu). Laser power output at the objective ranged from 20 to 100 mW, increasing with imaging depth. Images were acquired at a resolution of 794 × 527 pixels, corresponding to a physical area of 892 × 667 µm. To minimize contamination from ambient light, blackout material was used to shield the optical path from the visual display.

### 1-photon calcium imaging

Widefield 1-photon calcium imaging was used to map the retinotopic organization and delineate dorsal cortical area boundaries. Excitation light was provided by a 470 nm blue LED (Thorlabs), and fluorescence emission was collected through a 510/84 nm green emission filter (Semrock) using a CCD camera (PCO Edge 3.1). Imaging was acquired at 10 frames per second using a 2x objective lens (NA = 0.10, Thorlabs), yielding an image resolution of 1024 × 768 pixels across the field of view.

### Visual stimulation

Visual stimuli were presented on a calibrated 22-inch LCD monitor (Samsung 2233RZ, 1680 by 1050-pixel resolution, 60 Hz refresh rate), positioned 18 cm from the animal’s right eye. The screen covered 120° × 80° of the right visual field (0–120° azimuth; ±40° elevation). Custom software built in PsychoPy (Peirce, 2007) was used to control stimulus presentation and synchronize with imaging acquisition.

For retinotopic mapping, we presented a circular patch of flickering noise rotating anticlockwise along an elliptical trajectory around the display center (Supplementary Video S1). Each trial consisted of four rotations, followed by a 6s inter-trial interval, repeated across 30 trials.

For cellular calcium imaging, full-screen square wave drifting gratings of 12 directions and 6 combinations of spatial frequencies (0.04, 0.08, and 0.16 cpd) and temporal frequencies (1 and 4 Hz) were presented. Stimuli were presented in pseudo-randomized order at fixed 5s intervals with 3s of visual stimulation interleaved with 2s of equiluminant gray screen (54–80 cd/m²). In dark-condition experiments, the monitor and all external light sources (e.g., for pupil or facial tracking) were turned off.

### Viral vectors injections

To label neurons projecting to anterior and posterior RSC, retrograde AAV viral vectors were injected through a burr-hole craniotomy using a glass micropipette (30 µm tip diameter, Drummond Scientific Company, Pulling system from Sutter Instrument P-2000). Two combination viral vector with identical promotors were used: rgAAV-CAG-tdTomato with rgAAV-CAG-GFP, or rgAAV-hSyn-mCherry with rgAAV-hSyn-GFP (Table S3). Injections were targeted to the center of RSPd/v using stereotaxic coordinates relative to Bregma. For the anterior RSC, the coordinates were anteroposterior (AP) -1.5 mm, mediolateral (ML) 0.5 mm, and dorsoventral (DV) -0.8 mm. For the posterior RSC, the coordinates were AP -3.2 mm, ML 0.5 mm, and DV -0.8 mm. Each injection was made approximately 0.5 mm below the cortical surface. A volume of 150–200 nL viral solution was infused at each site, using a Nanoject II injector that delivered ∼13 nL per pulse at 20-second intervals.

### Histology

Three weeks following viral injection, mice were deeply anesthetized with an intraperitoneal injection of ketamine and xylazine (100 mg/kg and 10 mg/kg, respectively), and transcardially perfused first with phosphate-buffered saline (PBS), followed by 4% paraformaldehyde (PFA). Brains were extracted and cut into 100 µm-thick coronal sections using either a vibratome (VT1000 S, Leica) or a cryostat (CM3050 S, Leica). Sections were mounted using Vectashield containing the nuclear stain DAPI (4’,6-diamidino-2-phenylindole; Vector Laboratories). Confocal imaging was performed with a Zeiss LSM 900 microscope equipped with a 10x Plan-APOCHROMAT objective (NA = 0.45). Images were acquired using a 0.7x zoom setting with approximately 12 by 7 tile scans and a 15% overlap between tiles at a resolution of either 1.25 (1024 x 1024) or 2.5 µm (512 x 512) per pixel. Z-stacks were acquired with a step size of 5 - 7.5 µm and covered at least 80 µm in depth.

## QUANTIFICATION AND STATISTICAL ANALYSIS

All data analyses were performed using custom scripts written in Python and MATLAB.

### Widefield calcium imaging data

For Widefield calcium imaging, fluorescence signals were first normalized to the pre-stimulation baseline. A min-max filter was applied to enhance spatial and temporal contrast (Supplementary Video S1). Trial-averaged data were analyzed using a discrete Fourier transform along the temporal axis to extract pixel-wise frequency components, with amplitude representing response strength and phase indicating response timing. These components were visualized using an HSV colormap based on normalized amplitude and phase. Registration to the Allen Common Coordinate Framework (CCF) dorsal cortex was achieved by matching retinotopic maps in primary and higher visual areas (Figure S3).

### Cellular calcium imaging data

Cellular calcium imaging data were acquired using ScanBox and processed with Suite2P for motion registration, ROI detection, and calcium signal extraction (Pachitariu et al., 2016). The extracted neuronal activity included the raw fluorescence signal F_cell_ and neuropil fluorescence signal F_np_ . For each ROI, the raw somatic fluorescence signal was corrected by subtracting surrounding neuropil activity using the formula:

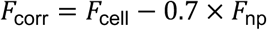

The baseline fluorescence was estimated using a min/max filter with 60s kernel size, applied to F_corr_, followed by Gaussian smoothing (SD = 10 time bins). Spike deconvolution was performed using the OASIS algorithm, which models the calcium trace as the convolution of spike events with an exponential decay kernel. A fixed decay constant (1.5s) appropriate for GCaMP6s was used (Friedrich et al., 2017).

### Selection of active neurons

To identify neurons actively engaged during the behavioral task, we applied separate criteria for movement-related and visually evoked activity. For task-related activity, lap reliability was calculated as the proportion of running laps in which a neuron exhibited significant calcium transients. Transients were considered significant if the fluorescence signal exceeded the baseline ΔF/F₀ by more than three times the standard deviation. This criterion ensured inclusion of neurons that consistently responded across repeated laps.

For assessing visual responsiveness, trial-averaged ΔF/F₀ traces for each stimulus condition were baseline-corrected by subtracting the median pre-stimulus signal, calculated over the five frames preceding stimulus onset. To quantify the consistency of visually evoked responses, a visual reliability index ()) was computed as the 75th percentile of cross-trial Pearson correlation coefficients of the de-randomized response time courses. A neuron was classified as visually responsive if) > 0.3 (Han et al., 2022).

A neuron was considered active during the task if it exhibited either lap reliability greater than 0.3 (i.e., significant activity in at least 30% of laps) or visual reliability greater than 0.3.

### Population decoding of position

We employed a Bayesian algorithm (Zhang et al., 1998; Davidson et al., 2009) to estimate the probability of position given the ΔF/F_0_ of a fixed number of randomly selected neurons. Bayesian decoding was performed while the animal was in a running epoch, defined as consecutive frames of forward movement lasting at least 1s, with a minimum speed of 5 cm/s. Consecutive epochs separated by less than 0.5s were merged (Danielson et al., 2016).

To compare the decoding error between the anterior and posterior retrosplenial cortex, odd trials were used for model training, while even-numbered trials were reserved for testing. To assess decoding error in darkness, 80% of the total trials from the light-on session were used for training, while the remaining trials were reserved for testing using 5-fold cross-validation. The Bayesian decoding equation is given by:

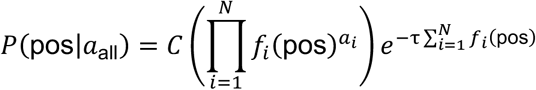

where 1_all_ represents the activity (ΔF/F_0_) of the neurons, and 2 is the normalization constant ensuring that the posterior sums to 1. The parameter 8 is the time bin size, which depends on the imaging sampling rate. The function 5_!_(pos)^"!^ represents the position-binned activity (with 1.5 cm per bin), and 9 is the number of neurons. The decoded position for each time bin was determined as the position with the maximum posterior probability. The decoding error was computed as the absolute difference between the actual position and the decoded position. Chance-level decoding was set at 50 spatial bins (75 cm)

### Detection of neurons with position activity

Position-tuned neurons were identified using a lower-bound threshold computed from deconvolved calcium data (Mao et al., 2020). The linear track was divided into 100 position bins (1.5 cm per bin), and the occupancy-normalized activity was smoothed using a Gaussian window (SD = 3 position bins) for each neuron.

To determine statistical significance, neuronal activity was circularly shifted by a random time between 20s and the total session duration minus 20s, repeated 200 times to generate a shuffled distribution. A neuron was classified as position-tuned if the lower bound of its actual activity (mean-SEM) in any position bin across trials exceeded the 97.5 percentile of the shuffled distribution.

Position-tuned neurons were further examined by quantifying their position-tuning properties. The position tuning field width was calculated from the number of consecutive position bins in which the mean activity exceeded 30% of the difference between peak and baseline activity (below 15 cm or above 120 cm were discarded). The neuron was considered a position cell if it met the following criteria: (1) Position tuning field width was a continuous region spanning at least 15 cm but no more than 120 cm. (2) At least one position tuning field was presented in more than one-third of all trials.

### Quantification of position-related activity

Spatial information (SI) was calculated using the following formula (Skaggs et al., 1993):

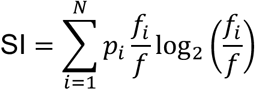

where *p_i_* is the occupancy probability (the fraction of the time spent in the -th position bin), 5_!_ is the occupancy-normalized deconvolved calcium activity (summed activity divided by the total time spent in the -th position bin), and 5 is the overall mean activity.

To examine activity reliability across the trials (whether exhibited calcium transients at the same locations, Figure 2F), we computed trial-to-trial correlation as the median of the pairwise Pearson correlation coefficients across trials using deconvolved calcium activity (Pettit et al., 2022).

### Characterization of direction and orientation visual selectivity

Orientation and direction tuning were quantified using the orientation selectivity index (OSI) and directional selectivity index (DSI), respectively:

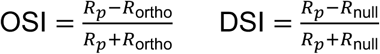

where A_/_is the response to the preferred orientation/direction, A_ortho_ is the response to the orthogonal orientation, A_null_ is the response to the 180° direction from the preferred direction. Neurons with OSI >0.33 or DSI >0.33 were considered orientation and direction-selective, respectively (Ohki et al., 2005).

### Quantification of retrogradely labeled cells

Labeled neurons were manually selected in different channels separately. Registration was performed using the DAPI channel across all brain slices to the Allen Brain Atlas (Shamash et al., 2018) (adapted from Allen Common Coordinate Framework, https://github.com/cortex-lab/allenCCF). After applying a transformation matrix to the brain slice, each ROI was registered to a specific brain region based on the mouse brain atlas. ROIs were classified according to the hierarchy structure tree provided by the Allen Brain Institute (Wang et al., 2020). A 3D whole-brain reconstruction with labeled ROIs was generated based on the model adapted from brainrender (Claudi et al., 2021) (https://github.com/brainglobe/brainrender).

To assess the relative input distribution between posterior and anterior subregions of the retrosplenial cortex (RSC), we computed a bias index defined as the log-ratio of the fractional input to posterior RSC versus anterior RSC:

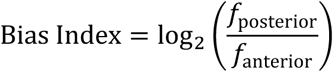

where 5_posterior_and 5_anterior_ represent the fraction of input to the posterior and anterior RSC, respectively. Positive values indicate a relative bias toward posterior RSC, whereas negative values indicate a bias toward anterior RSC (Figure 7H).

### Statistical testing

Statistical analyses were performed to assess differences in neuronal properties between anterior and posterior RSC across animals. To compare scalar summary measures such as the fraction of position-tuned or visually responsive neurons, position-tuning properties, decoding error, and spatiotemporal visual tuning metrics, we used the Mann–Whitney U test, a non-parametric alternative to the t-test that does not assume normality. For population-level comparisons of full distributions—such as cumulative distributions of position tuning properties, visual reliability, orientation selectivity, and direction selectivity, we used the two-sample Kolmogorov–Smirnov test, which evaluates differences in both the shape and central tendency of distributions. Statistical significance levels are reported as follows: p < 0.05 (*), p < 0.01 (**), p < 0.001 (***).

## Acknowledgments

We thank Ta-Shun Su for assistance with the data analysis code, Ben Vermaercke, and Adrien Philippon for technical assistance with *in vivo* imaging and the behavioral setups. YT.W. was supported MOE–KU Leuven Taiwan scholarships. J.C. acknowledges support from FWO (Fellowship 1226320N). V.B. acknowledges support from FWO (grant G0C1220N). V.B. and F.K. acknowledge support from FWO (grant G077321N). We are also grateful to the members of the Bonin and Kloosterman labs for valuable discussions and feedback.

## Author contributions

Conceptualization: YT.W., V.B. and F.K.; Methodology: YT.W. and V.B.; Software: YT.W.; Investigation: YT.W. and V.B. and F.K.; Data Analysis: YT.W.; Visualization: YT.W. Writing— original draft: YT.W.; Writing—review & editing: YT.W., V.B. and F.K.; Funding acquisition: J.C. and V.B.; Supervision: V.B. and F.K.

## SUPPLEMENTAL INFORMATION

**Table S1:** Animal Information for imaging.

| MOUSE | AGE<sup>*1</sup><br>(weeks) | SEX | HEMI-<br>SPHERE | FOV<sup>*2</sup><br>AM (mm) | FOV<sup>*2</sup><br>PM (mm) | FOV<sup>*2</sup><br>AL (mm) | FOV<sup>*2</sup><br>PL (mm) | GENOTYPE<sup>*3</sup> | Recording<br>session<sup>*4</sup> |
| --- | --- | --- | --- | --- | --- | --- | --- | --- | --- |
| YW006 | 20 | Male | Left | A: 0, -1.04<br>P: -0.23, -2.74 | A: 0, -2.17<br>P: -0.23, -3.74 | A: -0.67, -1.17<br>P: -0.9, -2.74 | A: -0.67, -2.17<br>P: -0.9, -3.74 | Thy1 | A: 2 (depth)<br>P: 2 (depth) |
| YW008 | 20 | Male | Left | A: 0, -1.13<br>P: -0.28, -3.17 | A: 0, -2.13<br>P: -0.28, -4.17 | A: -0.67, -1.13<br>P: -0.9, -2.74 | A: -0.67, -2.13<br>P: -0.95, -4.17 | Thy1 | A: 2 (depth)<br>P: 2 (depth) |
| YW010 | 22 | Male | Left | A: 0, -1.22<br>P: -0.12, -2.33 | A: 0, -2.22<br>P: -0.12, -3.33 | A: -0.67, -1.22<br>P: -0.79, -2.33 | A: -0.67, -2.22<br>P: -0.79, -3.33 | Thy1 | A: 1<br>P: 1 |
| YW017 | 22 | Male | Left | A: -0.22, -1.21<br>P: -0.2, -2.32 | A: -0.22, -2.21<br>P: -0.2, -3.32 | A: -0.89, -1.21<br>P: -0.87, -2.32 | A: -0.89, -2.21<br>P: -0.87, -3.32 | Thy1 | A: 1<br>P: 1 |
| YW022 | 25 | Male | Left | A: -0.2, -1.02<br>P: 0, -2.61 | A: -0.2, -2.02<br>P: 0, -3.61 | A: -0.87, -1.02<br>P: -0.67, -2.61 | A: -0.87, -2.02<br>P: -0.67, -3.61 | CaMKII | A: 1 (ETL)<br>P: 1 (ETL) |
| YW032 | 20 | Male | Left | A: 0, -1.2<br>P: 0, -2.94 | A: 0, -2.38<br>P: 0, -3.94 | A: -0.67, -1.38<br>P: -0.67, -2.94 | A: -0.67, -2.38<br>P: -0.67, -3.94 | CaMKII | A: 1 (ETL)<br>P: 1 (ETL) |
| YW033 | 20 | Male | Left | A: 0, -1.2<br>P: -0.1, -2.68 | A: 0, -2.2<br>P: -0.1, -3.68 | A: -0.67, -1.2<br>P: -0.77, -2.68 | A: -0.67, -2.2<br>P: -0.77, -3.68 | CaMKII | A: 1 (ETL)<br>P: 1 (ETL) |
| YW048 | 20 | Male | Left | A: -0.05, -1.038<br>P: -0.13, -2.59 | A: -0, -2.14<br>P: 0, -3.67 | A: -0.78, -1.09<br>P: -0.83, -2.69 | A: -0.66, -2.19<br>P: -0.71, -3.76 | CaMKII | A: 1 (ETL)<br>P: 1 (ETL) |

**\*1.** Age for the imaging data collection
**\*2.** Field of view anchor coordinates: Imaging regions were anchored at specific coordinates relative to Bregma in anteromedial (AM), posteromedial (PM), anterolateral (AL), and posterolateral (PL) locations. The anterior RSC (A) and posterior RSC (P) were analyzed, with negative values indicating positions in the left hemisphere (x) or posterior relative to Bregma (y).
**\*3.** Thy1: Thy1-GCaMP6s; CaMKII: CaMKII-tTA x TRE-GCamp6.lineG6s2
**\*4.** Recording sessions: Two animals underwent two separate recording sessions, each corresponding to a different imaging depth, recorded on separate days.

**Table S2:**
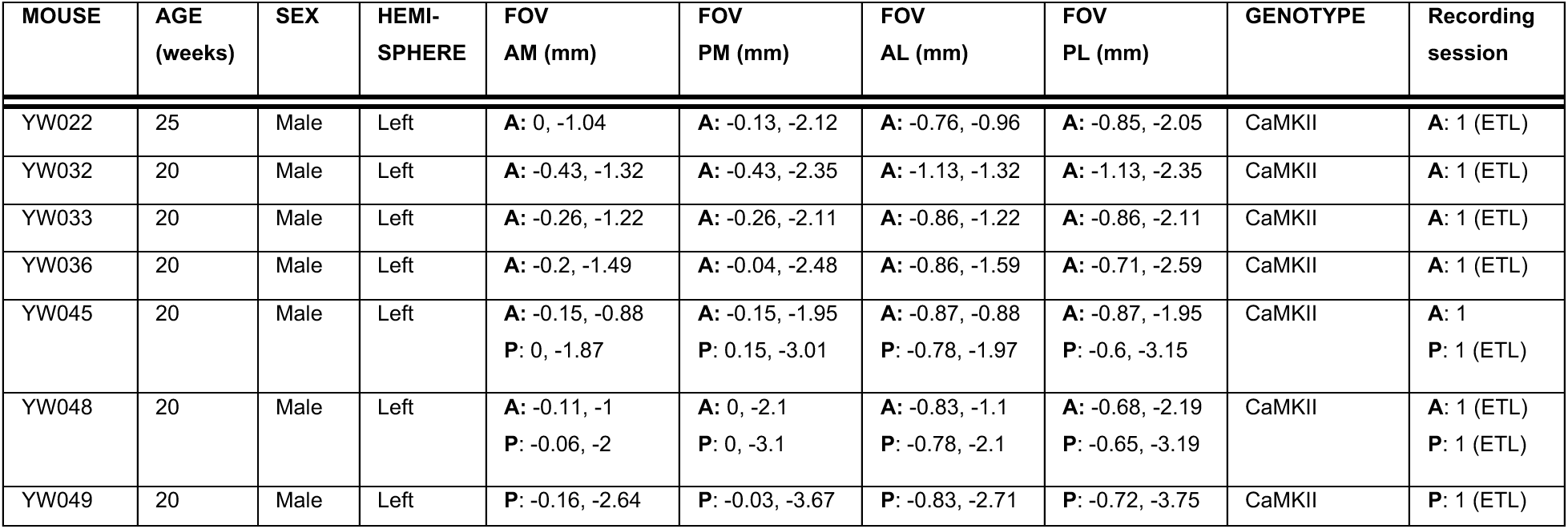
Animal information for darkness experiment.

| MOUSE | AGE<br>(weeks) | SEX | HEMI-<br>SPHERE | FOV<br>AM (mm) | FOV<br>PM (mm) | FOV<br>AL (mm) | FOV<br>PL (mm) | GENOTYPE | Recording<br>session |
| --- | --- | --- | --- | --- | --- | --- | --- | --- | --- |
| YW022 | 25 | Male | Left | A: 0, -1.04 | A: -0.13, -2.12 | A: -0.76, -0.96 | A: -0.85, -2.05 | CaMKII | A: 1 (ETL) |
| YW032 | 20 | Male | Left | A: -0.43, -1.32 | A: -0.43, -2.35 | A: -1.13, -1.32 | A: -1.13, -2.35 | CaMKII | A: 1 (ETL) |
| YW033 | 20 | Male | Left | A: -0.26, -1.22 | A: -0.26, -2.11 | A: -0.86, -1.22 | A: -0.86, -2.11 | CaMKII | A: 1 (ETL) |
| YW036 | 20 | Male | Left | A: -0.2, -1.49 | A: -0.04, -2.48 | A: -0.86, -1.59 | A: -0.71, -2.59 | CaMKII | A: 1 (ETL) |
| YW045 | 20 | Male | Left | A: -0.15, -0.88<br>P: 0, -1.87 | A: -0.15, -1.95<br>P: 0.15, -3.01 | A: -0.87, -0.88<br>P: -0.78, -1.97 | A: -0.87, -1.95<br>P: -0.6, -3.15 | CaMKII | A: 1<br>P: 1 (ETL) |
| YW048 | 20 | Male | Left | A: -0.11, -1<br>P: -0.06, -2 | A: 0, -2.1<br>P: 0, -3.1 | A: -0.83, -1.1<br>P: -0.78, -2.1 | A: -0.68, -2.19<br>P: -0.65, -3.19 | CaMKII | A: 1 (ETL)<br>P: 1 (ETL) |
| YW049 | 20 | Male | Left | P: -0.16, -2.64 | P: -0.03, -3.67 | P: -0.83, -2.71 | P: -0.72, -3.75 | CaMKII | P: 1 (ETL) |

**Table S3:**
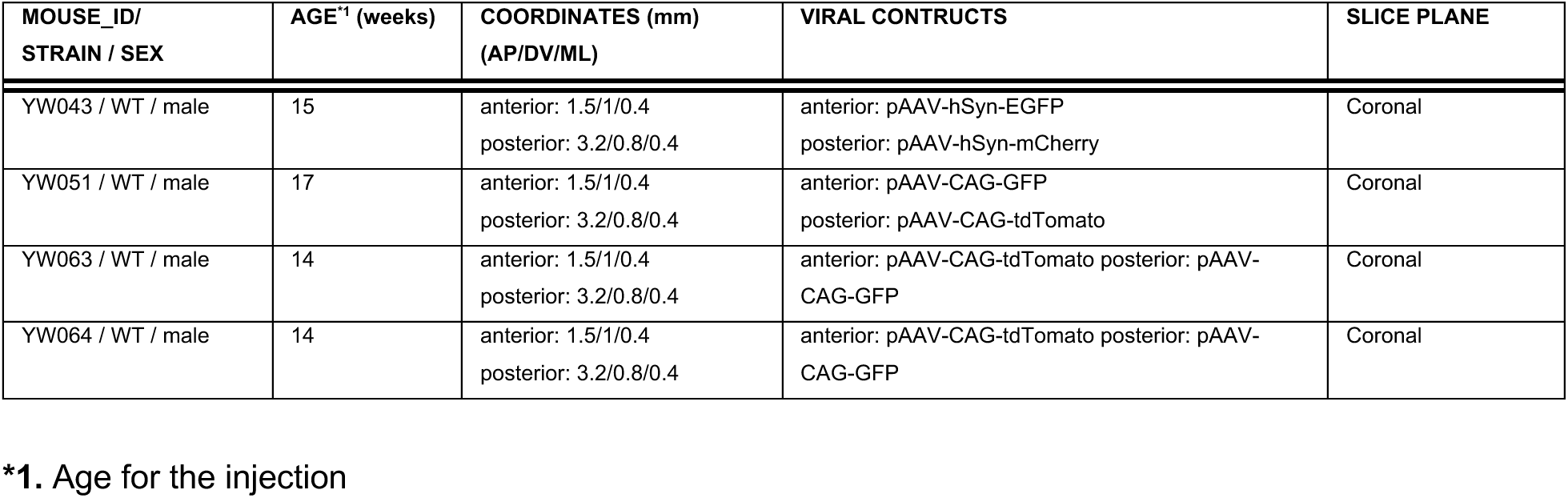
Animal information for retrogradely tracing.

| MOUSE_ID/<br>STRAIN / SEX | AGE<sup>*1</sup> (weeks) | COORDINATES (mm)<br>(AP/DV/ML) | VIRAL CONSTRUCTS | SLICE PLANE |
| --- | --- | --- | --- | --- |
| YW043 / WT / male | 15 | anterior: 1.5/1/0.4<br>posterior: 3.2/0.8/0.4 | anterior: pAAV-hSyn-EGFP<br>posterior: pAAV-hSyn-mCherry | Coronal |
| YW051 / WT / male | 17 | anterior: 1.5/1/0.4<br>posterior: 3.2/0.8/0.4 | anterior: pAAV-CAG-GFP<br>posterior: pAAV-CAG-tdTomato | Coronal |
| YW063 / WT / male | 14 | anterior: 1.5/1/0.4<br>posterior: 3.2/0.8/0.4 | anterior: pAAV-CAG-tdTomato posterior: pAAV-CAG-GFP | Coronal |
| YW064 / WT / male | 14 | anterior: 1.5/1/0.4<br>posterior: 3.2/0.8/0.4 | anterior: pAAV-CAG-tdTomato posterior: pAAV-CAG-GFP | Coronal |

**\*1. Age for the injection**

**Figure S1.**
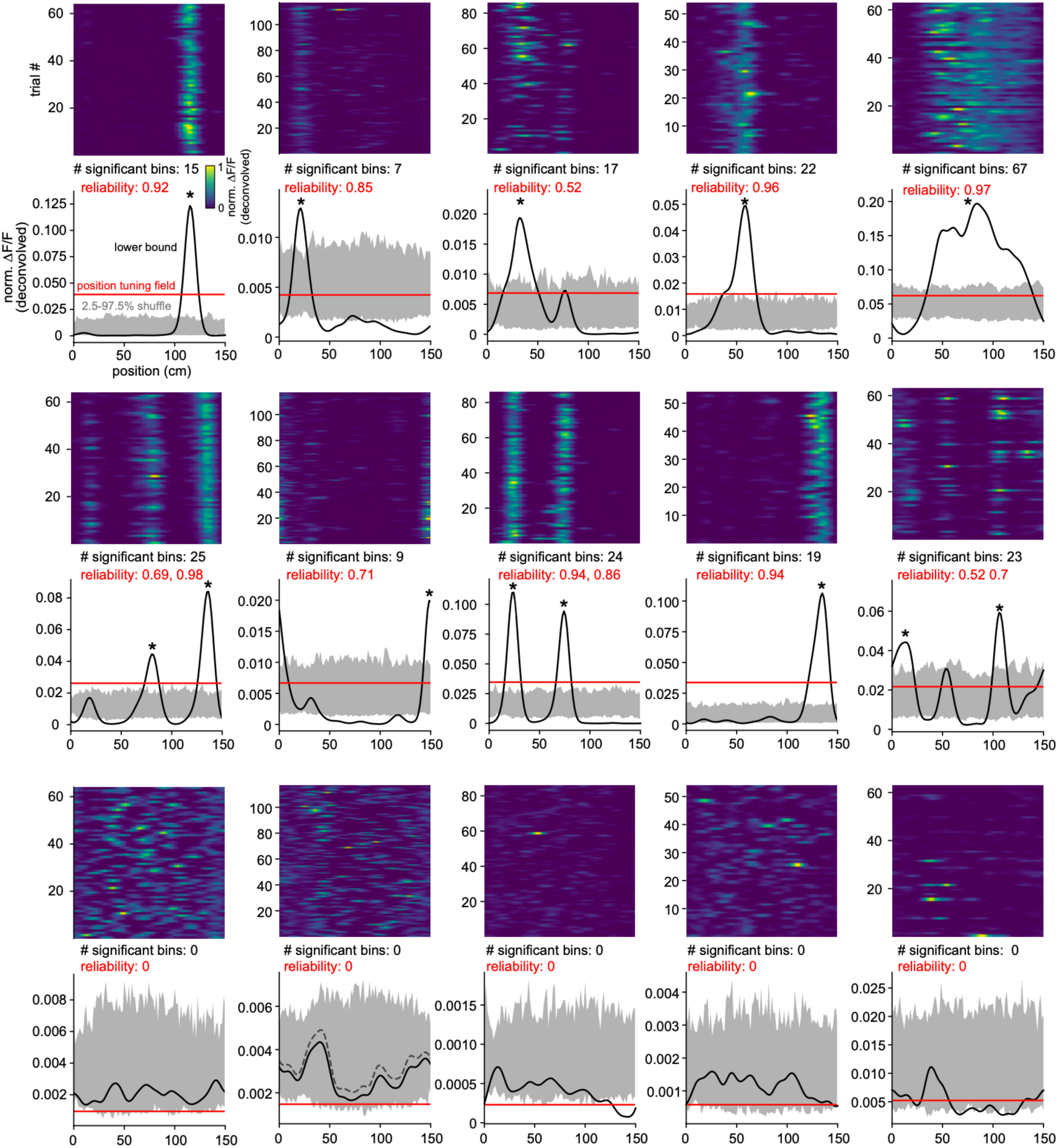
Selection of position-tuned neurons. Position-tuned neurons were identified based on the reliability of their position-related responses and further refined using of quantification of the position tuning field properties (see STAR Methods). The heatmaps show normalized, deconvolved ΔF/F0 signals from example RSC neurons, aligned to the animal’s position across repeated laps. Neurons were classified as position tuned if the lower bounds of their activity (mean – SEM) (black lines) exceeds the 97.5th percentile of the shuffled distribution (grey shaded area) in at least one spatial bin. In addition, neurons were required to have a position tuning field width between 15 cm and 120 cm (red line), and to exhibit reliable activation, defined as the presence of a detectable position tuning field in more than one-third of trials (reliability > 0.33). Neurons meeting these criteria are indicated with asterisks (*).

**Figure S2.**
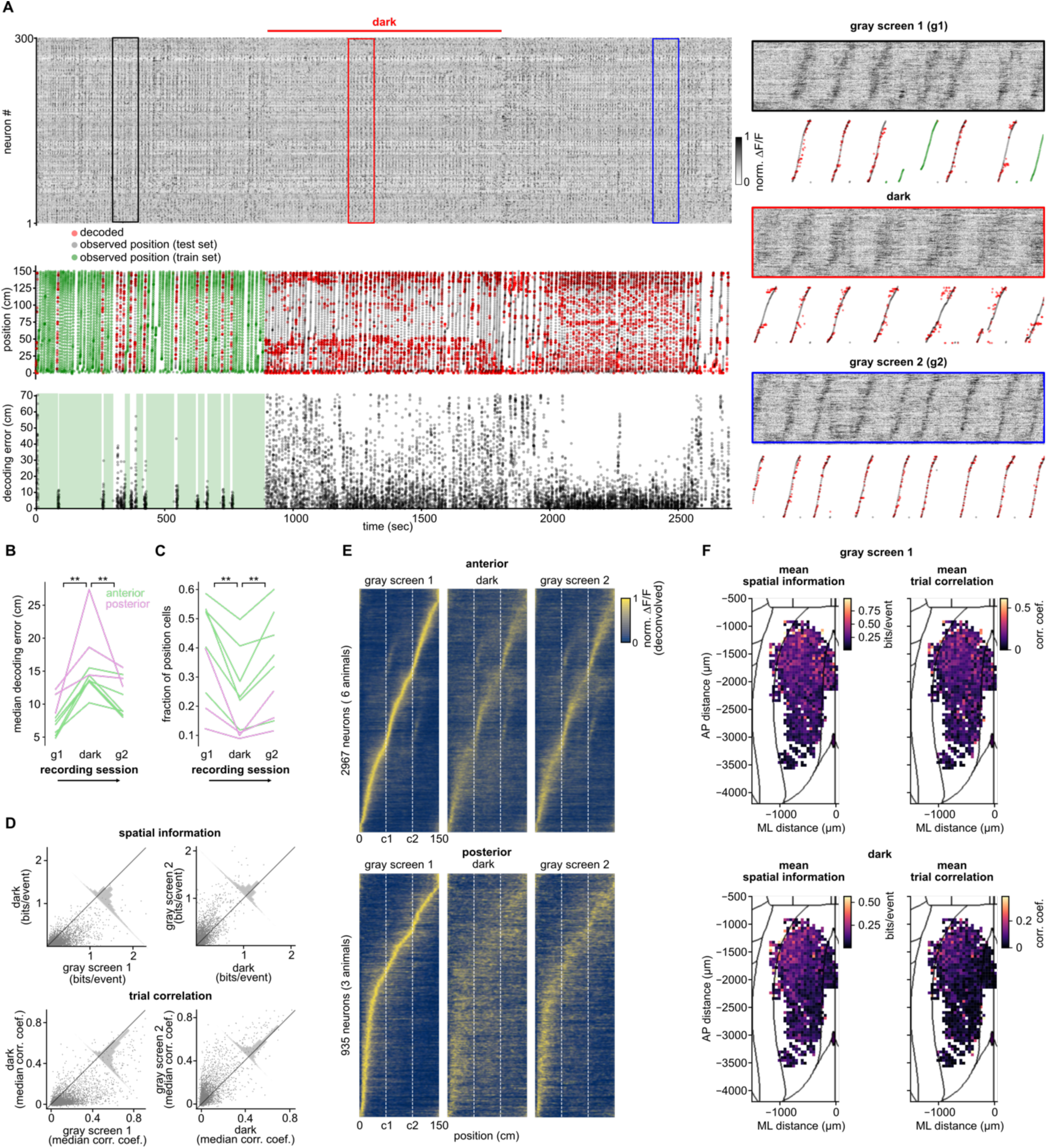
RSC sequences and position responses in darkness. (A) Top row: Example population activity from 300 randomly selected neurons, with magnified segments displayed on the right. Second row: a Bayesian decoder trained on 80% of trials from the initial gray screen session (green) and evaluated on the remaining trials using 5-fold cross-validation. Observed positions from test trials are indicated by gray dots, while decoded positions are represented by red dots. Bottom row: Frame-by-frame decoding error, with trained trials indicated by green shading. (B) Median decoding error across recording sessions. Wilcoxon signed-rank test; light-on 1 vs. dark; p = 0.0039; dark vs. light-on 2; p = 0.0039; n = 9 sessions from 7 animals. (C) Fraction of position-tuned neurons across sessions. Wilcoxon signed-rank test; gray screen 1 vs. dark; p = 0.0039; dark vs. gray screen 2; p = 0.0039; n = 9 sessions. (D) Pairwise scatter plots of spatial information and median trial-to-trial correlation across recording sessions, with histograms along the diagonal indicating the fraction of cells; n = 9 sessions. (E) Trial-averaged, deconvolved ΔF/F0 for all position-tuned neurons in anterior and posterior RSC across recording sessions, sorted according to activity from the initial gray screen session. (F) Dorsal cortical map showing mean spatial information and mean trial-to-trial correlation across the RSC. Median values were binned every 50 µm and represented using a color scale; n = 9 sessions.

**Figure S3.**
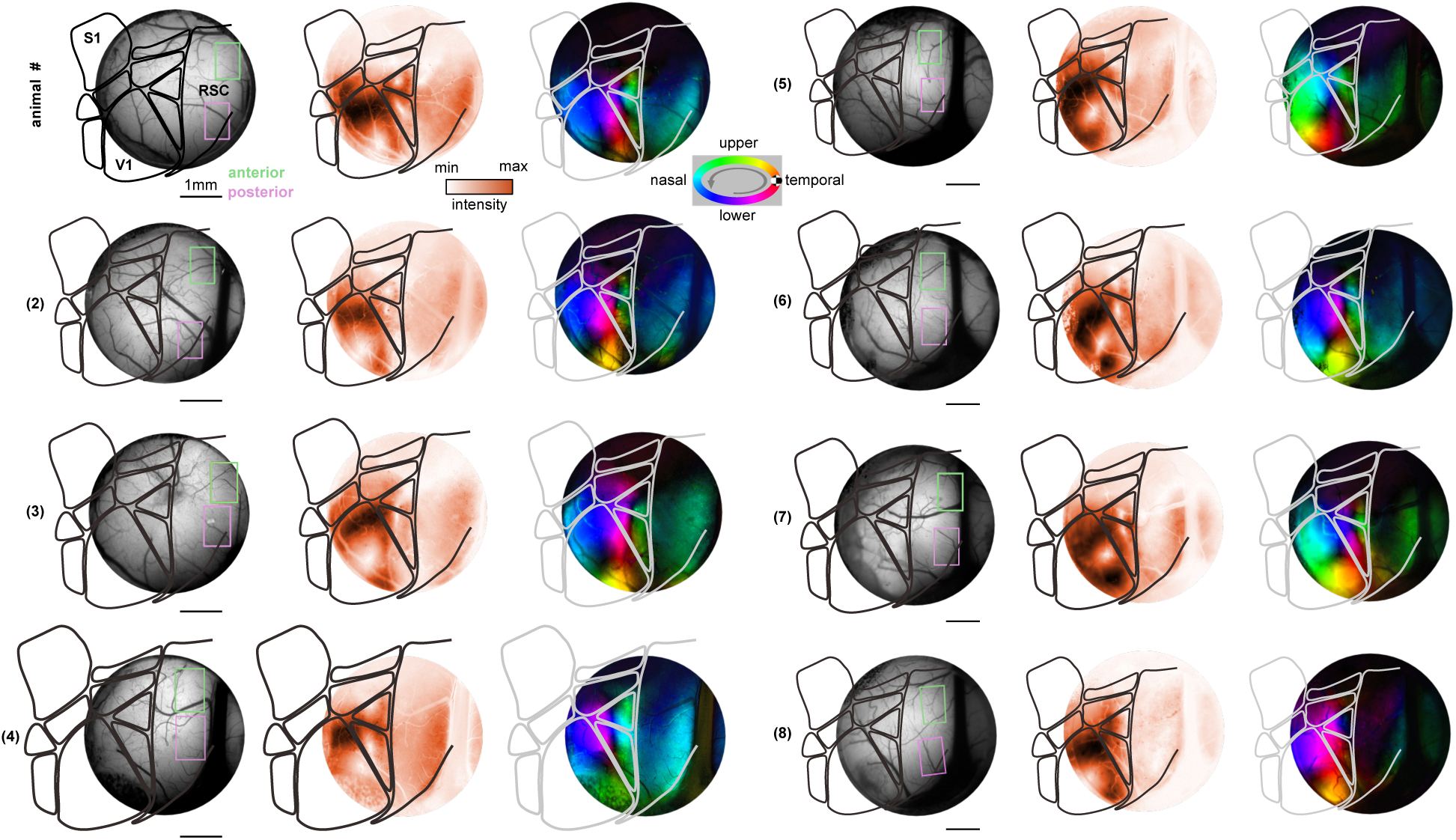
Selection of RSC imaging fields of view based on visual retinotopy. Imaging fields of view (FOVs) were selected from animals listed in Table S1. The left panels show a bright-field image of the cranial window, with outlines indicating inferred visual area borders (black) and the selected imaging FOVs (green and magenta). The middle panels show widefield calcium response amplitudes to a retinotopic stimulus (rotating circular patch), color-coded by magnitude. The right panels present the inferred retinotopic maps, where lightness encodes response magnitude and hue indicates the visual field location eliciting the strongest responses. Scale bar: 1 mm.

**Figure S4.**
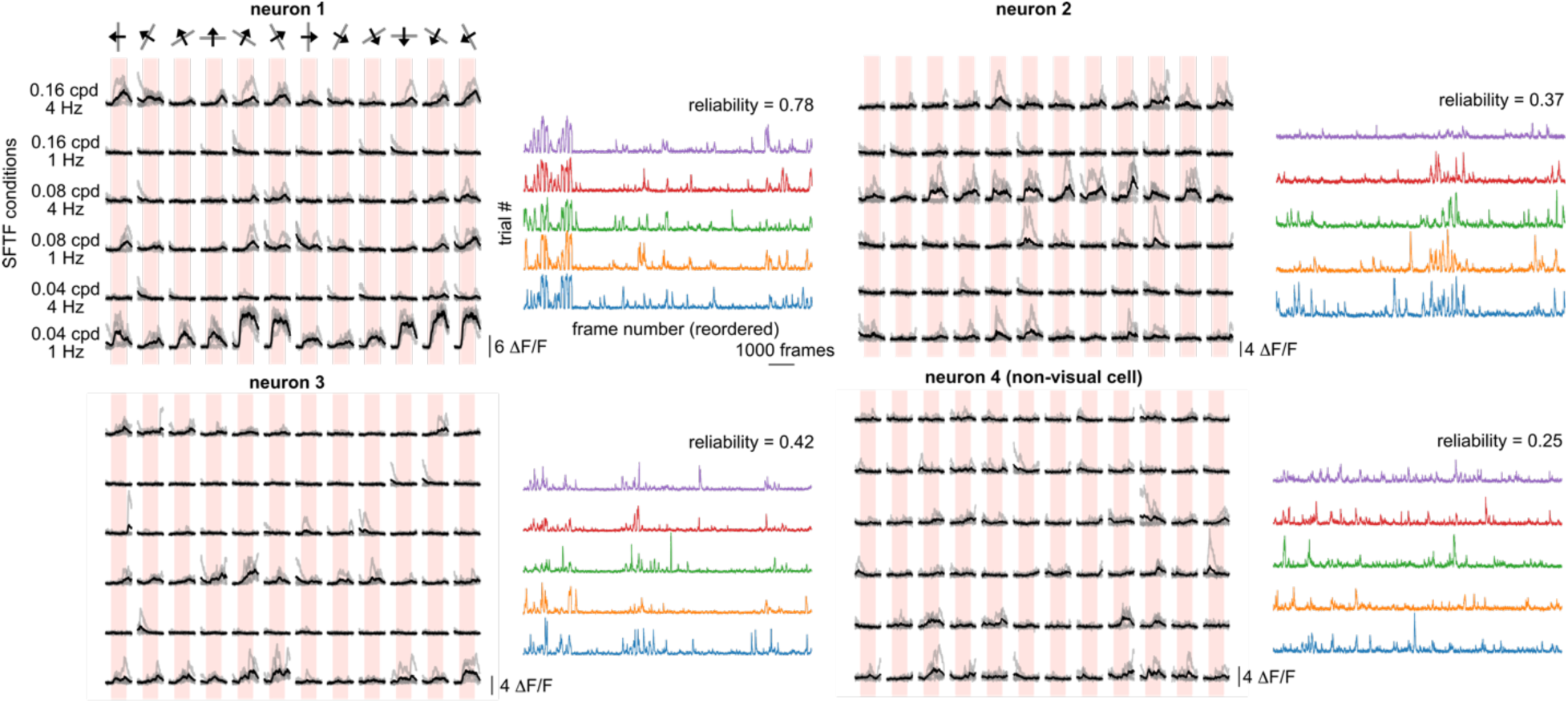
Selection of visually responsive cells. Four example RSC neurons responding to drifting grating visual stimuli. The left panel shows responses across different combinations of spatial frequencies (0.04, 0.08 and 0.16 cpd) and temporal frequencies (1 and 4 Hz), with rows representing frequency conditions and columns indicating stimulus direction. The right panel presents the corresponding calcium signals (ΔF/F0), de-randomized by visual stimulus conditions during stimulation epochs, with colors denoting individual trials. Reliability index ("), computed as the 75th percentile of trial-to-trial Pearson correlation coefficients (see STAR Methods), is indicated above each panel.

**Figure S5.**
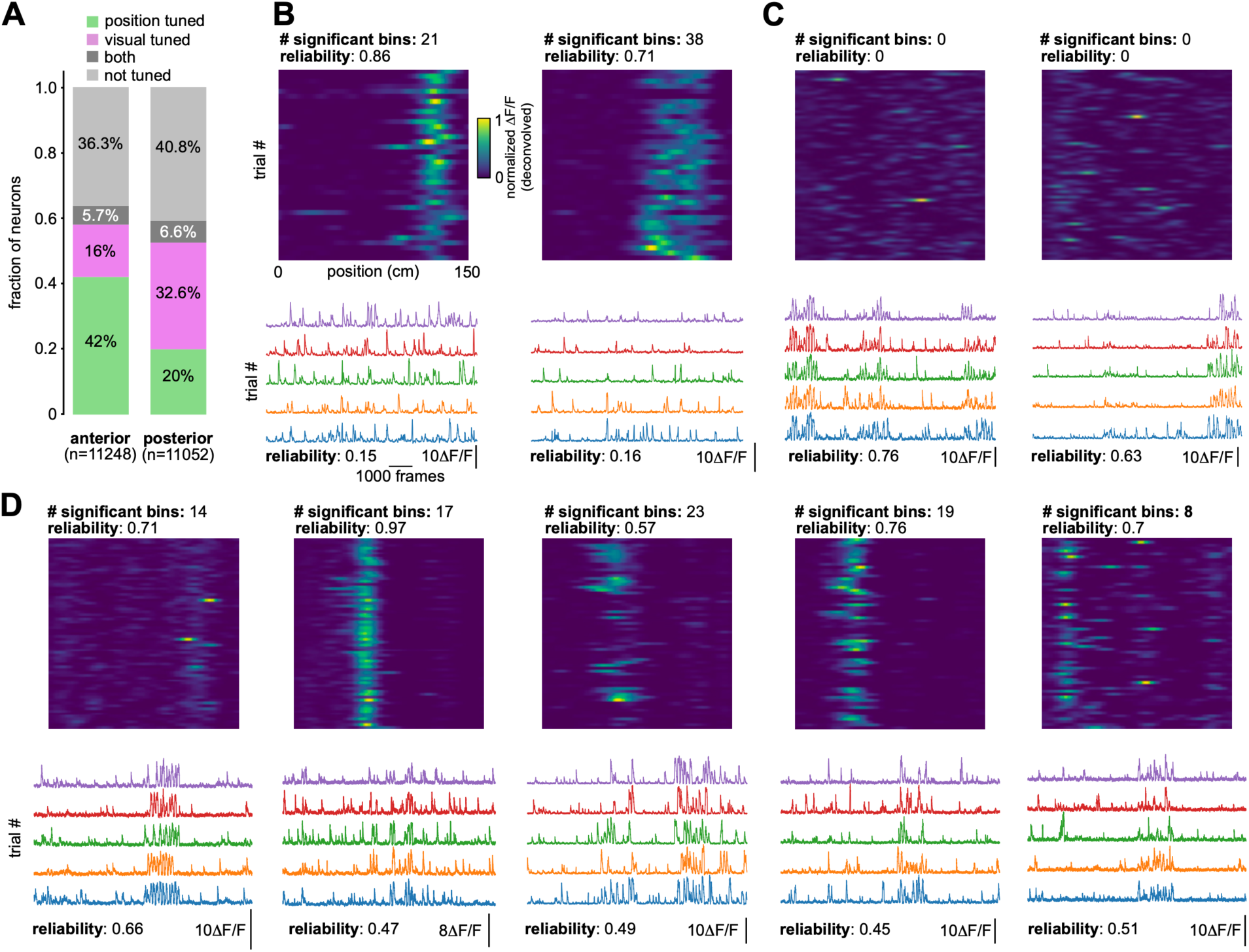
Multimodal responses in anterior and posterior RSC. (A) Proportion of neurons showing position tuned, visually responsive or bimodal responses in anterior and posterior RSC. (B–D) Examples of neuron responses: (B) Two position-tuned neurons. (C) Two visually responsive neurons. (D) Five neurons exhibiting both position tuning and visual responsiveness. Heat maps (top) show position related activity across repeated trials, while the traces show the de-randomized ΔF/F0 responses to visual stimulation, with different colors representing individual trials.

